# Wildlife in Cameroon harbor diverse coronaviruses including many isolates closely related to human coronavirus 229E

**DOI:** 10.1101/2021.09.03.458874

**Authors:** Nkom F. Ntumvi, Valantine Ngum Ndze, Amethyst Gillis, Joseph Le Doux Diffo, Ubald Tamoufe, Jean-Michel Takuo, Moctar M. M. Mouiche, Julius Nwobegahay, Matthew LeBreton, Anne W. Rimoin, Bradley S. Schneider, Corina Monagin, David J. McIver, Sanjit Roy, James A. Ayukekbong, Karen Saylors, Damien O. Joly, Nathan D. Wolfe, Edward M. Rubin, Christian E. Lange

## Abstract

Zoonotic spillover of animal viruses into human populations is a continuous and increasing public health risk. SARS-CoV-2 highlights the global impact emergence events can have. Considering the history and diversity of coronaviruses (CoVs), especially in bats, SARS-CoV-2 will likely not be the last to spillover from animals into human populations.

We sampled and tested wildlife in the central African country Cameroon to determine which CoVs are circulating and how they relate to previously detected human and animal CoVs. We collected animal and ecological data at sampling locations and used family-level consensus PCR combined with amplicon sequencing for virus detection.

Between 2003 and 2018, samples were collected from 6,580 animals of several different orders. CoV RNA was detected in 175 bats, a civet, and a shrew. The CoV RNAs detected in the bats represented 17 different genetic clusters, coinciding with alpha (n=8) and beta (n=9) CoVs. Sequences resembling human CoV-229E (HCoV-229E) were found in 40 *Hipposideridae* bats. Phylogenetic analyses place the human derived HCoV-229E isolates closest to those from camels in terms of the S and N genes, but closest to isolates from bats for the E, M, and RdRp genes. The CoV RNA positivity rate in bats varied significantly (p<0.001) between the wet (8.2%) and dry season (4.5%). Most sampled species accordingly had a wet season high and dry season low, while for some the opposite was found.

Eight of the suspected CoV species of which we detected RNA appear to be entirely novel CoV species, which suggests that CoV diversity in African wildlife is still rather poorly understood. The detection of multiple different variants of HCoV-229E-like viruses supports the bat reservoir hypothesis for this virus, with the phylogenetic results casting some doubt on camels as an intermediate host. The findings also support the previously proposed influence of ecological factors on CoV circulation, indicating a high level of underlying complexity to the viral ecology. These results indicate the importance of investing in surveillance activities among wild animals to detect all potential threats as well as sentinel surveillance among exposed humans to determine emerging threats.

## Introduction

The emergence of Severe Acute Respiratory Syndrome Coronavirus 2 (SARS-CoV-2) has highlighted the inherent risks and potential consequences of pathogen spillovers from animal reservoirs into the human populations. Animals are known to be the reservoir for many human diseases historically and contemporarily, as exemplified by zoonotic diseases such as rabies, brucellosis, bubonic plague, and trichinellosis [Shope 1982; Weiss 2003; Wolfe 2007; Gottstein 2009; Drancourt 2016; Cross 2019]. Humans are often considered an accidental host for zoonotic diseases in this context, even if human-to-human transmission is possible. This understanding has shifted with the advent, and increased availability, of genetic characterization of pathogens, and humans are now often considered opportunistic rather than accidental hosts. This is an especially apt description for viruses, since we now know that a significant number of pathogens commonly referred to as “human viruses” did not originally evolve with humans, but spilled over from animals more recently and subsequently adapted to humans [Weiss 2003; Wolfe 2007]. The best example for this may be HIV, which originated from multiple non-human primate spillover events in Africa during the early 20^th^ century, and it also applies to measles virus, influenza A viruses, and others [Weiss 2003; Sharp 2011]. While these and SARS-CoV-2 are some of the most publicized examples, there is a general trend of increasing infectious diseases outbreaks in humans, in particular of viral and zoonotic agents, over the past decades, potentially due to factors such as land use and climate change, population growth, and increased international trade and mobility [Jones 2008; Smith 2014; Allen 2017].

With the global SARS-CoV-2 pandemic, we appear to be witnessing such a post-spillover adaptation in real time, and there is strong evidence that this is not the first time an animal coronavirus (CoV) has gone through this process. While primary attention has previously focused on SARS-CoV, Middle Eastern Respiratory Syndrome (MERS)-CoV, and closely related CoVs, it has become clear that the four CoVs that are associated with the common cold (HCoV-NL63, HCoV-229E, HCoV-OC43, HCoV-HKU1), all likely derived from animal CoVs in the previous decades to centuries [Drosten 2003; Kan 2005; Yip 2009; Drexler 2019; Zaki 2012; Huynh 2012; Corman 2014; Corman 2015; Forni 2017; Corman 2018; Cui 2019]. Unlike SARS-CoV-1 and MERS-CoV, these four established themselves in the human population in a process that may have been similar to the current SARS-CoV-2 pandemic, albeit likely much slower. Genetic analysis suggests that the two alpha coronaviruses HCoV-NL63 and HCoV-229E originated in bats, like SARS-CoVs and MERS-CoV, while the beta CoVs HCoV-OC43 and

HCoV-HKU1 likely originated in rodents; in either case with or without a potential intermediate host [Pfefferle 2009; Tao 2017; Zhou 2020]. The fact that we know of three CoVs that spilled over into humans in just the past two decades, and that spillovers of the other four can be traced back to recent centuries suggests that these events are common, especially since it is estimated that only a minority of spillover events lead to continued transmission and detection in humans [Glennon 2019; Letko 2020]. CoV spillovers will thus likely continue to occur in the future. This indicates that a better understanding of animal CoVs will be useful to determine spillover risks and the biological mechanisms and drivers for diversification and spillover, and to develop appropriate prevention, mitigation, and treatment strategies for future CoV spillovers.

It is generally accepted that there is a direct link between a close genetic relationship of host species and the likelihood of interspecies transmission, however considering the biology of influenza A viruses, for example, where we see spillover from pigs and birds into humans, or CoVs where bats, rodents, camels, cows, and civets may play a role, it is by no means a hard rule [Parrish 2008; Kan 2005; Dennehy 2017]. With less closely related host species, the risks are more difficult to determine, but much emphasis has recently been placed on host diversity as an indicator for viral diversity, and the role of contact rates [Wolfe 2005; Pike 2010; Maganga 2014; Anthony 2017; Dennehy 2017; Leopardi 2018; Ntumvi 2021]. Host diversity likely results in more viruses circulating in a biome, and also provides more opportunities for interspecies transmission, host plasticity, and viral recombination [Dennehy 2017]. Host plasticity in animals may in turn be a predictor for a virus’ ability to be transmitted from human to human, and hence a major risk factor [Kreuder Johnson 2015]. Consequently, regions with a high biodiversity, such as large parts of Central Africa, South America, and Southeast Asia may be considered hot spots for spillover [Jones 2008; Allen 2017]. Reports from several African countries suggest there are many CoVs circulating, primarily in bats, including species related to pathogens such as SARS-CoV-1, MERS-CoV, HCoV-229E, and HCoV-NL63 [Tong 2009; Tao 2012; Annan 2013; Geldenhuys 2013; Razanajatovo 2015; Anthony 2017; Tao 2017; Geldenhuys 2018; Markotter 2019; Nziza 2019; Maganga 2020; Kumakamba 2021].

While the viral genome provides a lot of information about viruses, their origins, and their hosts, other factors also need to be considered when evaluating risks. Direct or indirect human-animal interaction is a prerequisite for zoonotic transmission, but human and animal behaviors and ecologies can play key roles in this process that may involve multiple steps of interspecies transmission and adaptation [Wolfe 2005; Wolfe 2007; Pike 2010; Maganga 2014; Euren 2020; Kumakamba 2021; Ntumvi 2021]. Previous studies have for example identified a high host plasticity for certain bat CoVs, indicating that these may potentially pose a higher zoonotic risk than those primarily adapted to a single host [Anthony 2017]. Other factors influencing transmission may include the type of human-animal interface and seasonal fluctuations of CoV circulation, as observed in bat populations [Montecino-Latorre 2020; Kumakamba 2021; Grange 2021]. Though important, data on many of these factors is still limited and needs further exploration. Studying CoV diversity in Africa promises rich data that could improve our understanding of their biology, evolutionary history and risks for humans.

The Central African country Cameroon, which includes some of the northern part of the Congo Basin, where HIV is believed to have spilled over into humans, is rich in biodiversity, with wildlife interaction being common for a large part of the rural population. Increased bushmeat trade and diverse wildlife living in close proximity with human populations makes some of these areas hotspots for high-risk interfaces between animals and humans [Wolfe 2005; Mickleburgh 2009; Saylors 2021].

Our goal was to determine what CoVs are circulating in wild animals, including rodents, bats, and non-human primates (NHPs), and assess if key ecological factors may influence the rate of CoV detection, and thus the exposure risk.

## Materials and Methods

### Sample and field data collection

Sample acquisition methods differed depending on the species and interface. Animals in peridomestic settings were captured and released after sampling (bats, rodents and shrews only), while samples from the (bushmeat) value chain were collected from freshly killed animals voluntarily provided by local hunters upon their return to the village following hunting, or by vendors at markets. Non-invasive fecal samples were collected from free-ranging NHPs, while some NHP samples, such as blood or serum, were collected during routine veterinary exams in zoos and wildlife sanctuaries. To avoid incentivizing hunting, hunters and vendors were not compensated. Identification was done in the field by trained field ecologists, as well as retrospectively based on field guides and other resources including those by Kingdon and Monadjem [Kingdon 2005; Monadjem 2010; Monadjem 2015]. Oral and rectal swab samples were collected into individual 2.0 ml screw-top cryotubes containing 1.5 ml of either Universal Viral Transport Medium (BD), RNA later, lysis buffer, or Trizol® (Invitrogen), while pea-sized tissue samples were placed in 1.5ml screw-top cryotubes containing 500ul of either RNA later or lysis buffer (Qiagen), or without medium. Specimen collection was approved by an Institutional Animal Care and Use Committee (IACUC) of UC Davis (#16067, # 17803, and # 19300), JHU (ACR #2007-110-03, #2007-110-02 and #2007-110-11) and UCLA (Protocols # FS03M221 and FS06H205) and the Government of Cameroon. All samples were stored in liquid nitrogen as soon as practical, before being stored long-term in freezers at −80°C. Sample collection staff were trained in safe collection techniques in collaboration with representatives from Ministry of Fisheries Livestock Animal Industries (MINEPIA), the Ministry of Forestry and Wildlife (MINFOF), and the Ministry of Environment Nature Protection and Sustainable Development MINEPDED) and wore dedicated clothing, N95 masks, nitrile gloves, and protective eyewear during animal capture, handling, and sampling.

### Sample processing

All laboratory work was carried out at Cameroon’s Military Health Research Center (CRESAR). RNA was extracted either manually using Trizol®, with an Qiagen AllPrep kit (tissue), Qiagen Viral RNA Mini Kit (swabs collected prior to 2014), or with a Zymo Direct-zol RNA kit (swabs collected after 2014) and stored at −80°C. Afterwards RNA was converted into cDNA using a GoScript™ Reverse Transcription kit (Promega), and stored at −20°C until analysis. Two conventional nested broad range PCR assays, both targeting conserved regions within the RNA-Dependent RNA Polymerase gene (RdRp) were used to screen samples for coronavirus RNA. The first PCR amplifies a product of approximately 286 nucleotides between the primer binding sites, and was specifically designed for the detection of a broad range of coronaviruses [Quan 2010]. The second PCR was used in two modified versions, with one of them specifically targeting a broad range of coronaviruses in bats and the second one broadly targeting coronaviruses of other hosts [Watanabe 2010]. Both versions amplify 387 nucleotides between the primer binding sites. Followup PCRs were designed to amplify the 3’-prime end of the Spike (S), as well as the Envelope (E), Membrane (M) and Nucleoprotein (N) genes of CoVs related to HCoV-229E (Table 1).

**Table 1:**
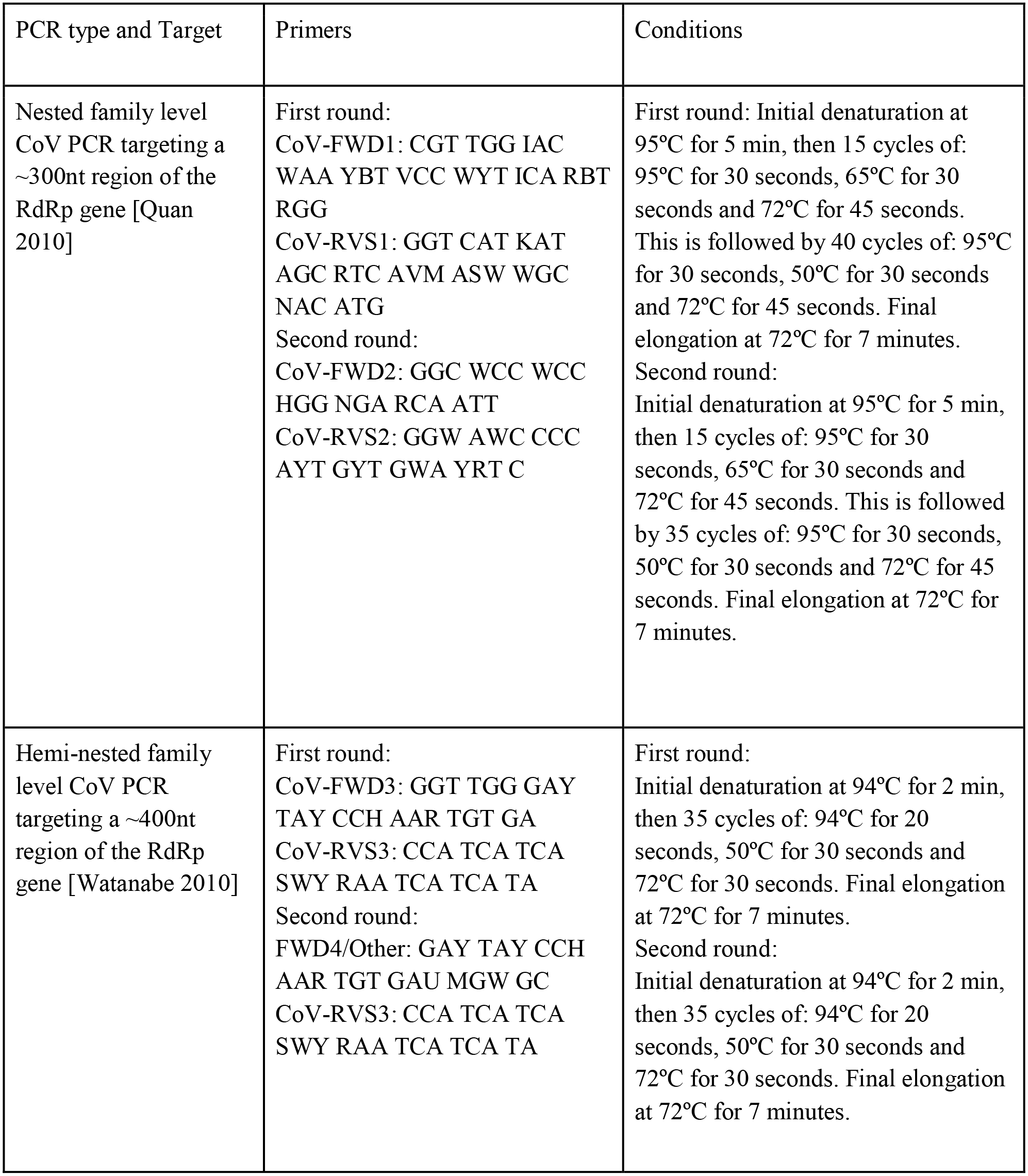
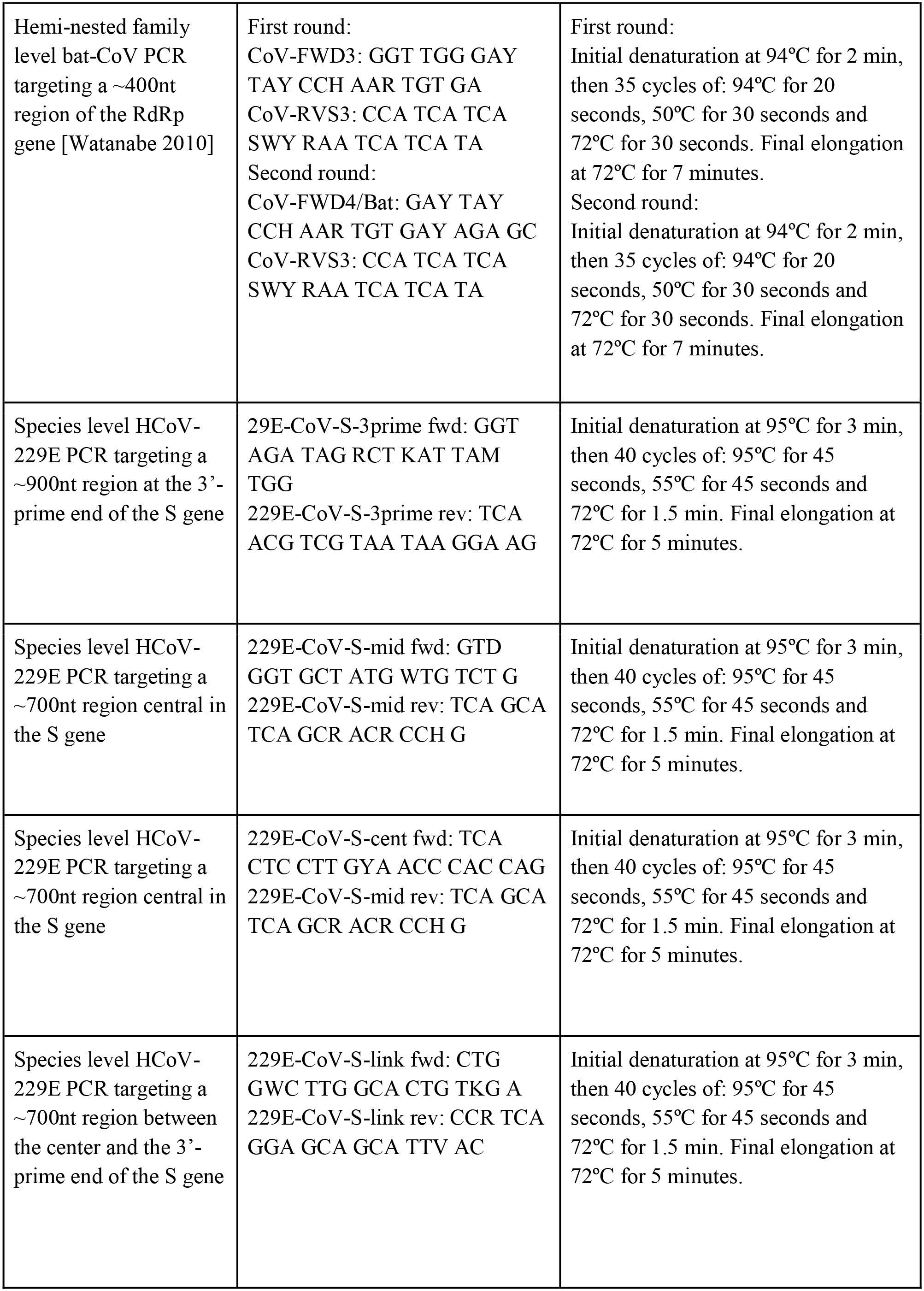
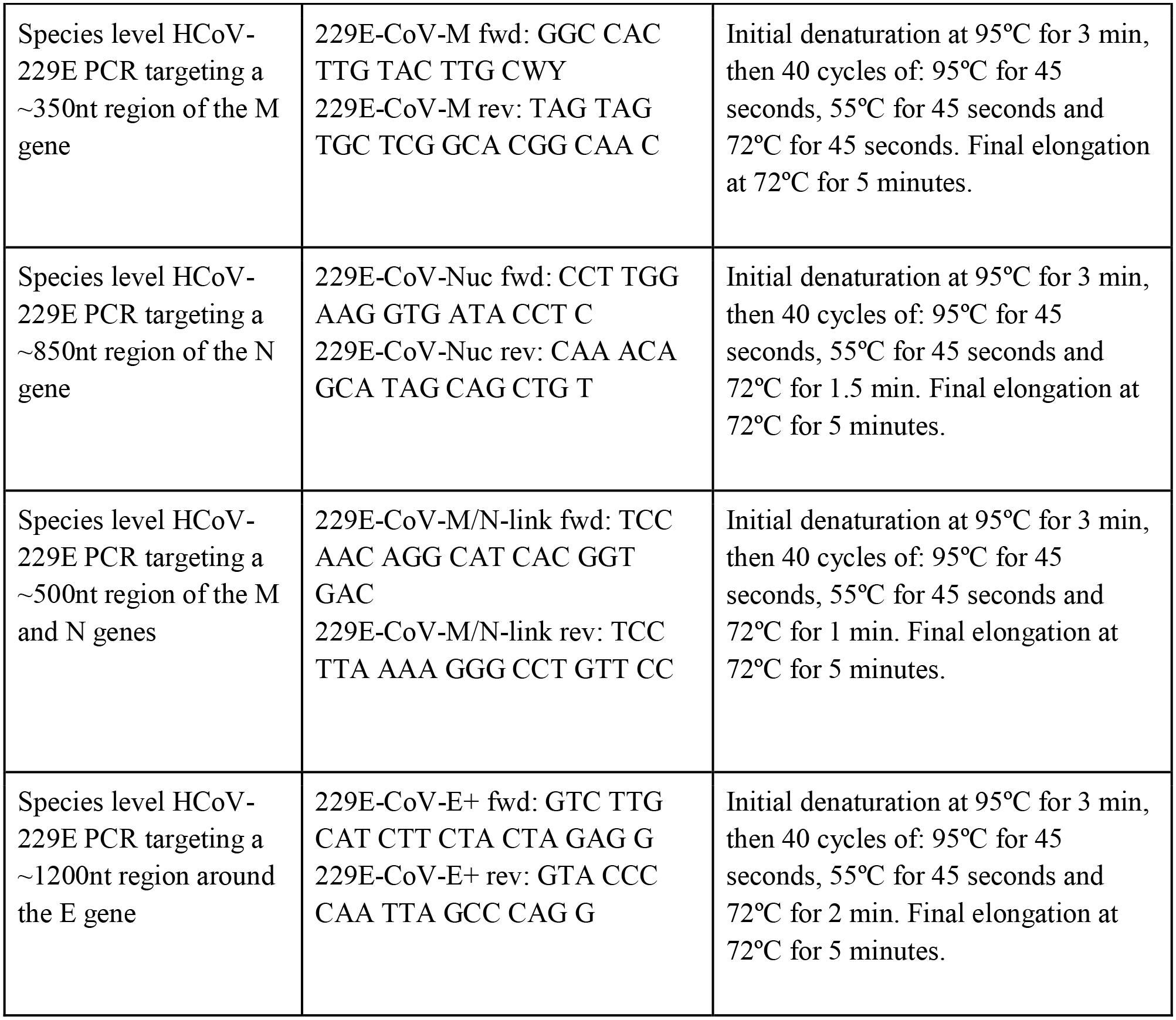
PCR primers and protocols.

PCR products were subjected to gel electrophoresis on a 1.5% agarose gel to identify amplicons of the expected size. PCR was repeated for all samples where this was the case, and PCR products were excised. Amplified DNA was extracted using either the QIAquick Gel Extraction Kit (Qiagen) or the Wizard SV Gel and PCR Cleanup System (Promega) and sent for commercial Sanger sequencing at either GATC, First Base or Macrogen. Extracts with low DNA concentrations were cloned prior to sequencing. All results from sequencing were analyzed in the Geneious 7.1 software, and primer trimmed consensus sequences compared to the GenBank database (BLAST N, NCBI). Samples where the PCR results could not be repeated, or where the sequencing did not yield interpretable sequences consistent with a CoV were counted as CoV RNA negative. All viral sequences obtained were deposited in the GenBank (Table 2, Supplement 1).

**Table 2:**
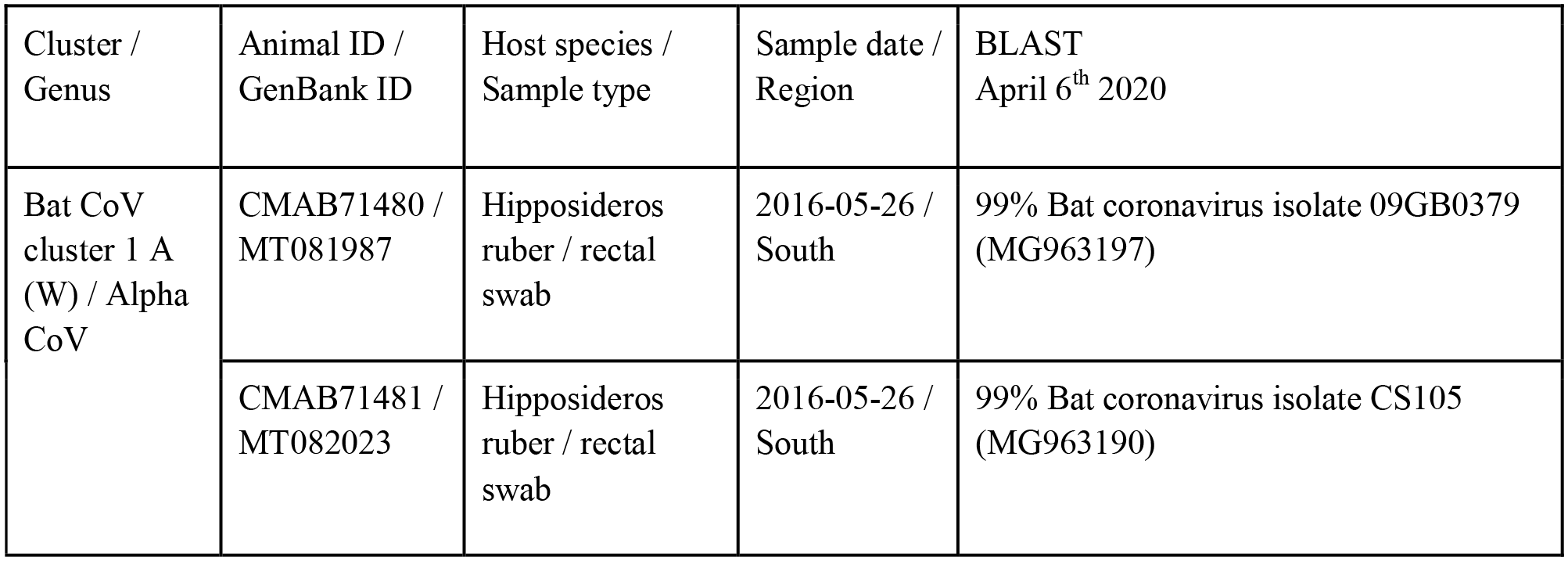
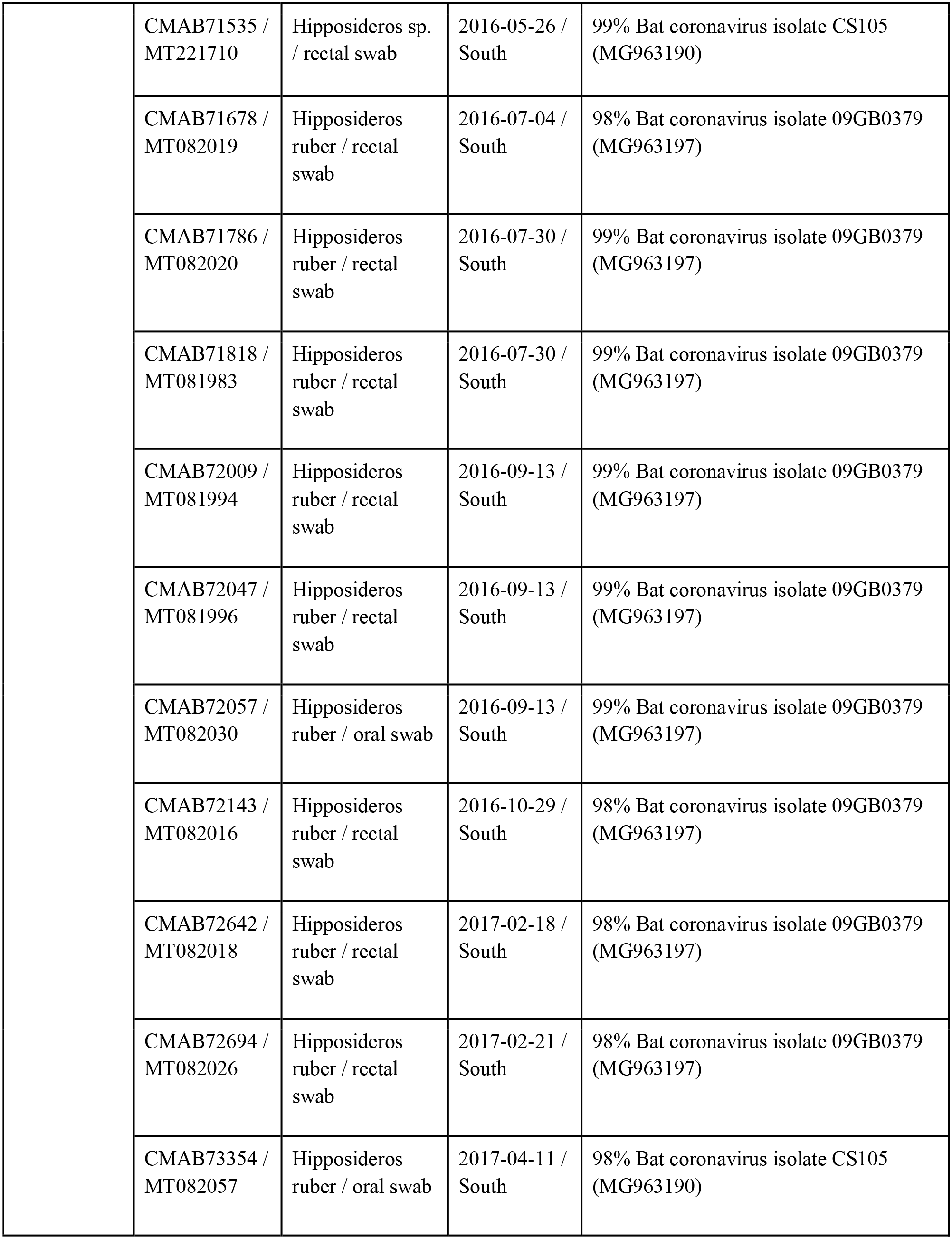
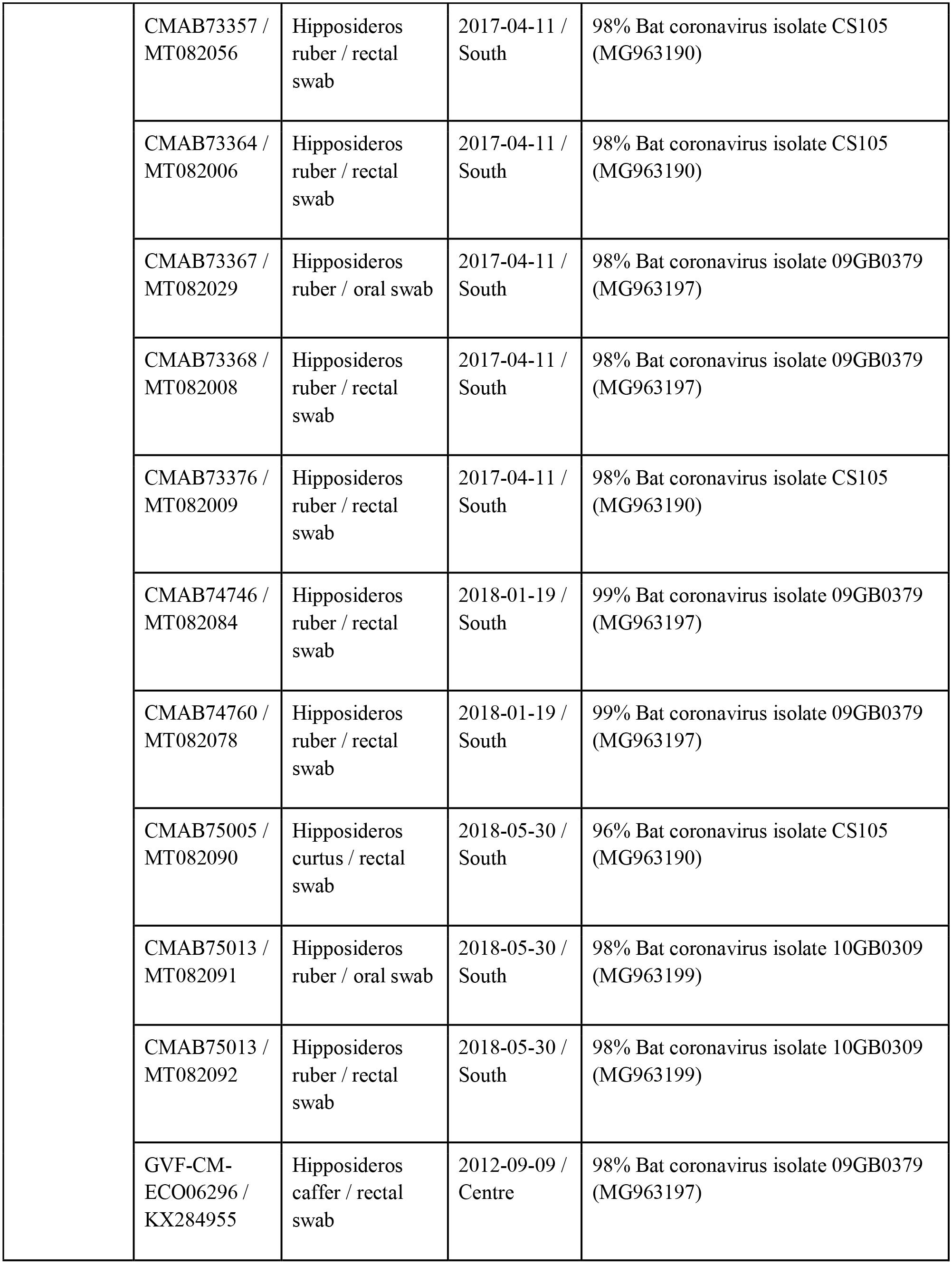
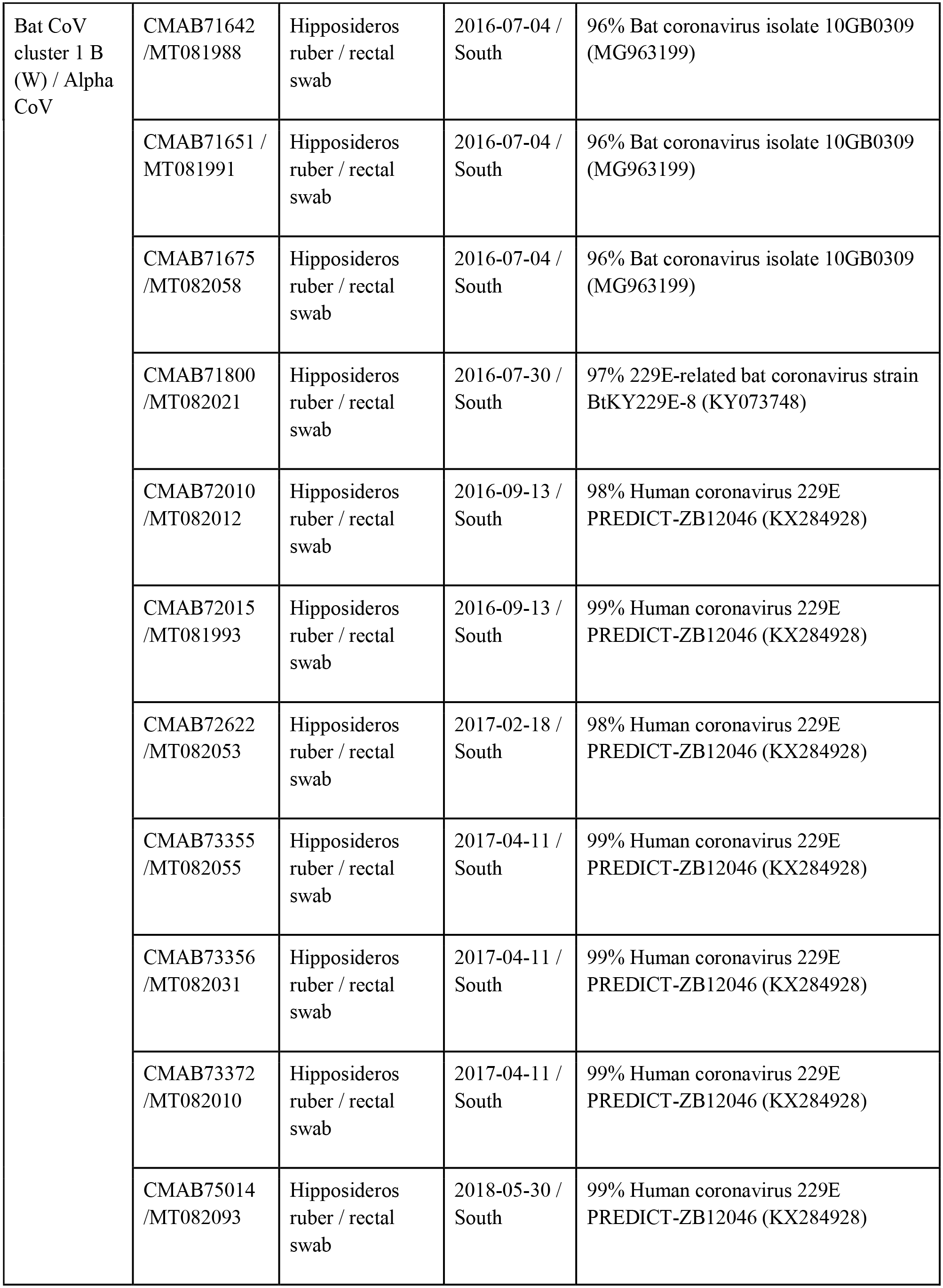
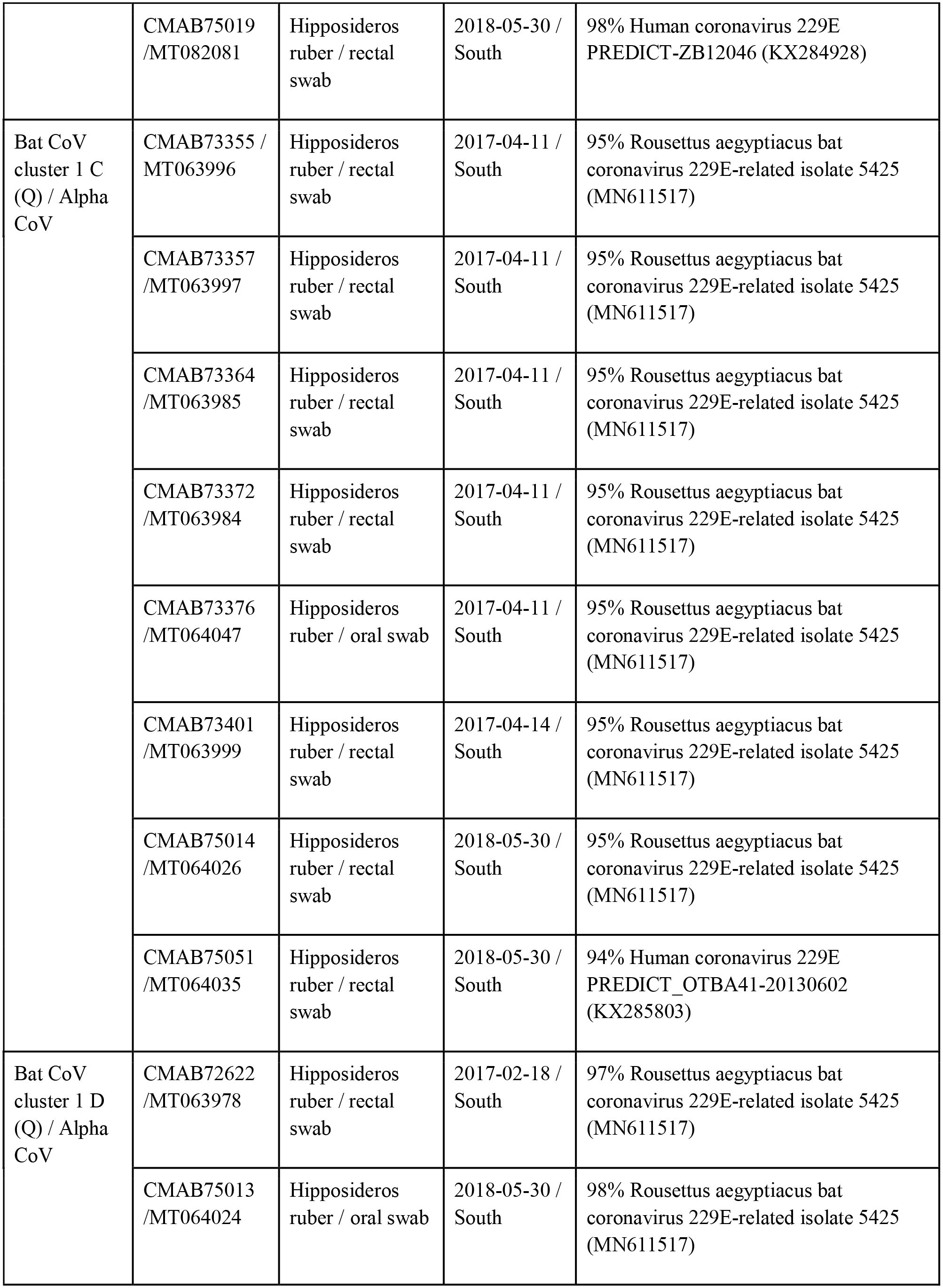
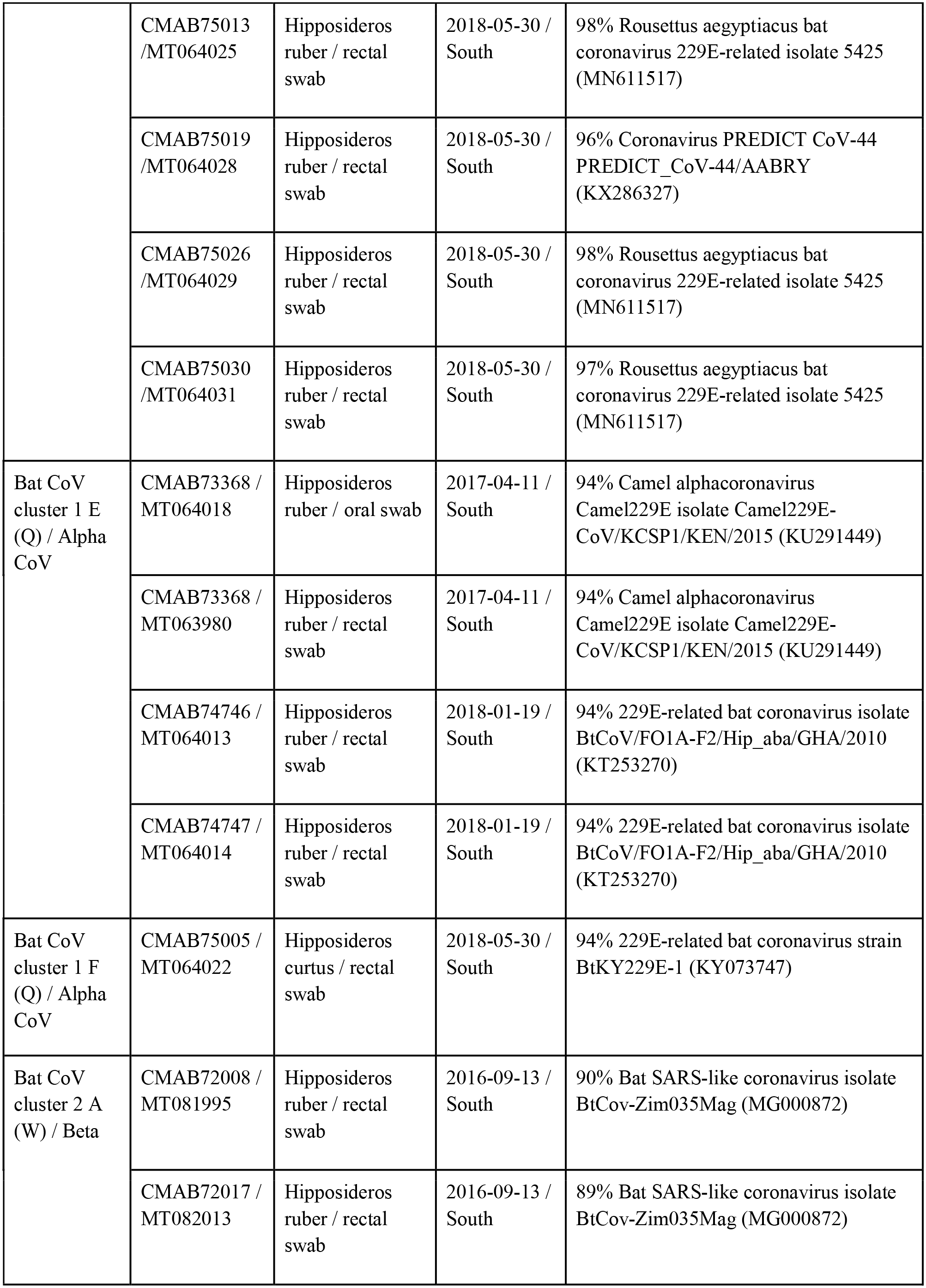
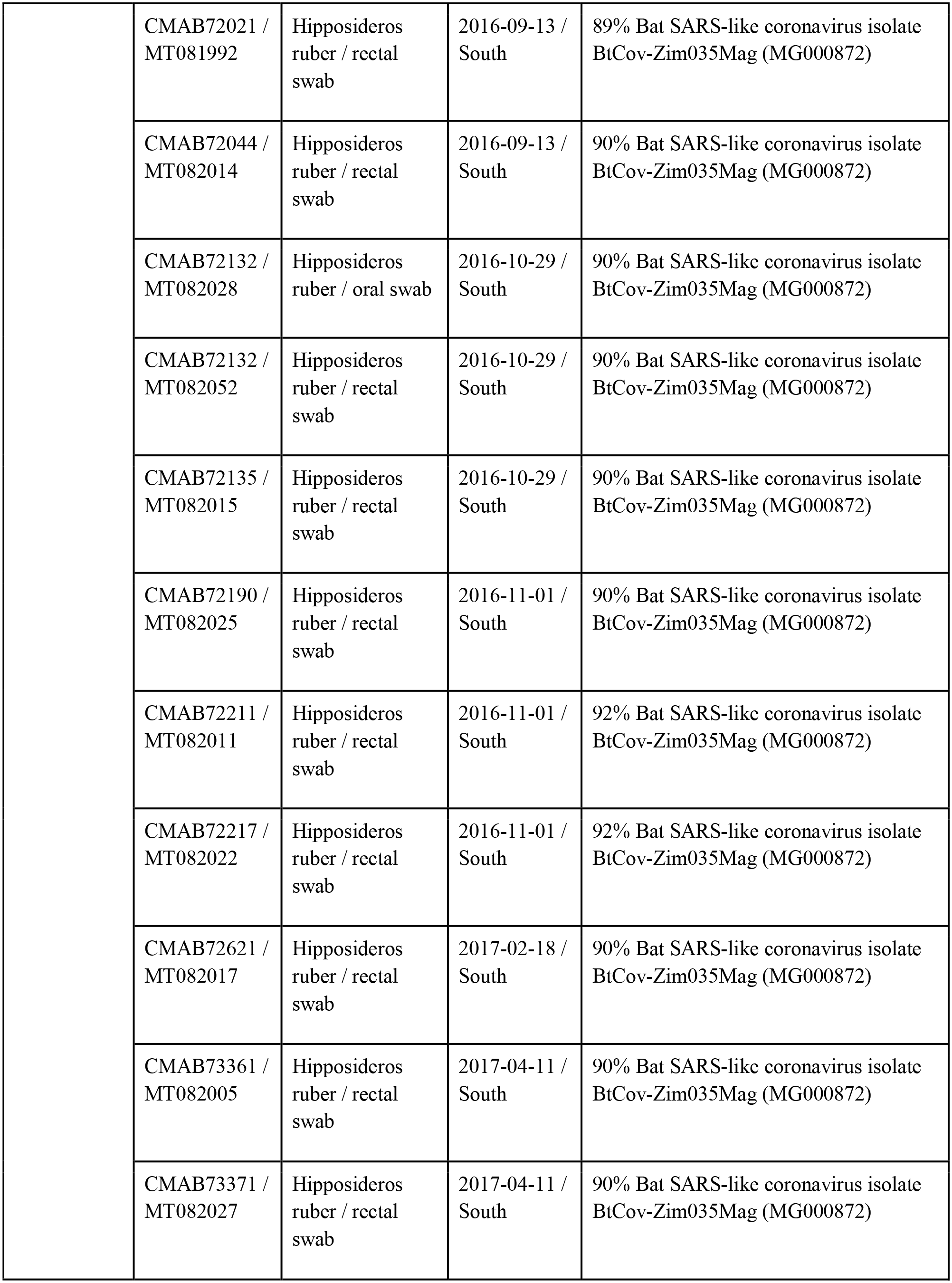
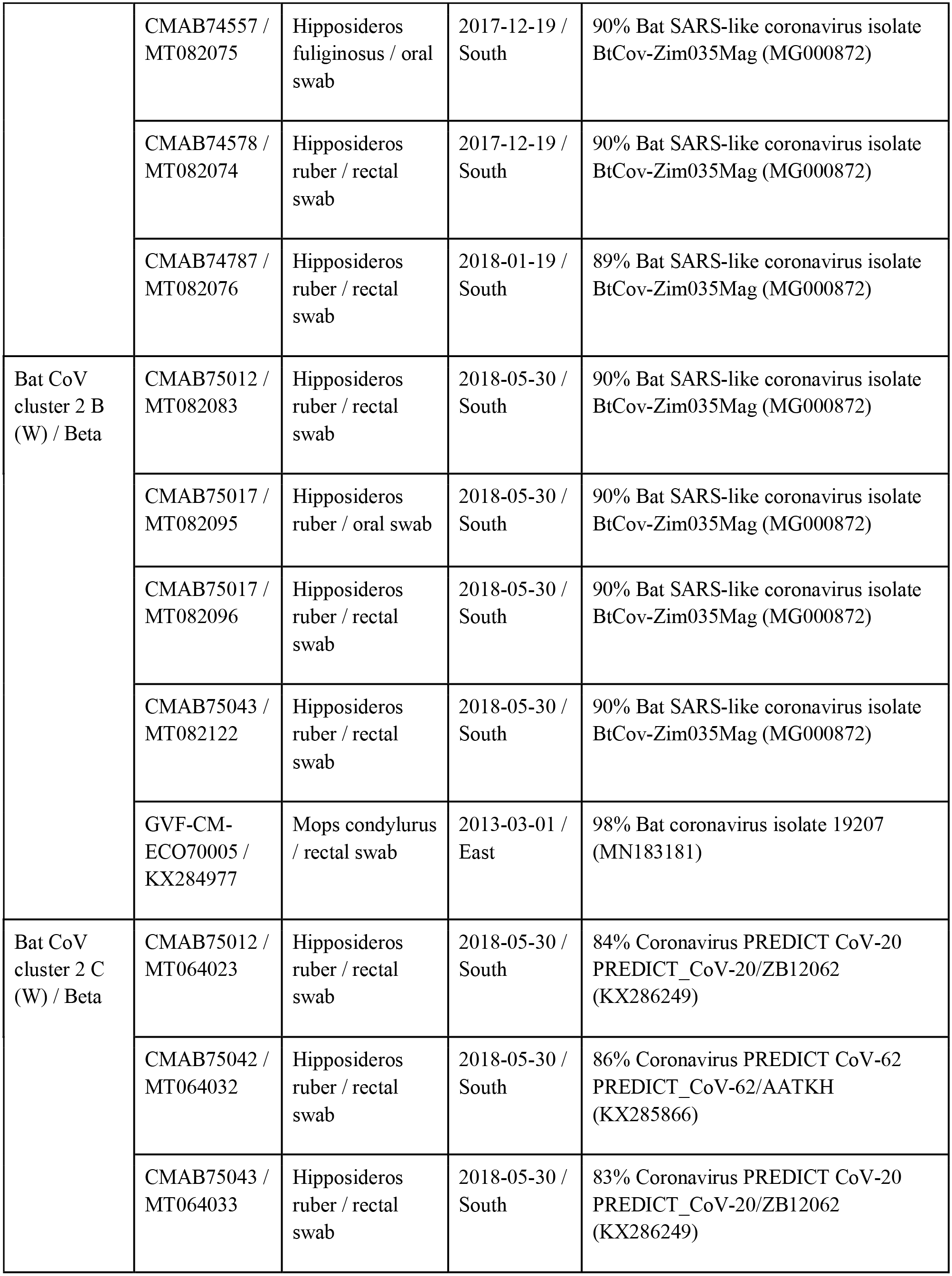
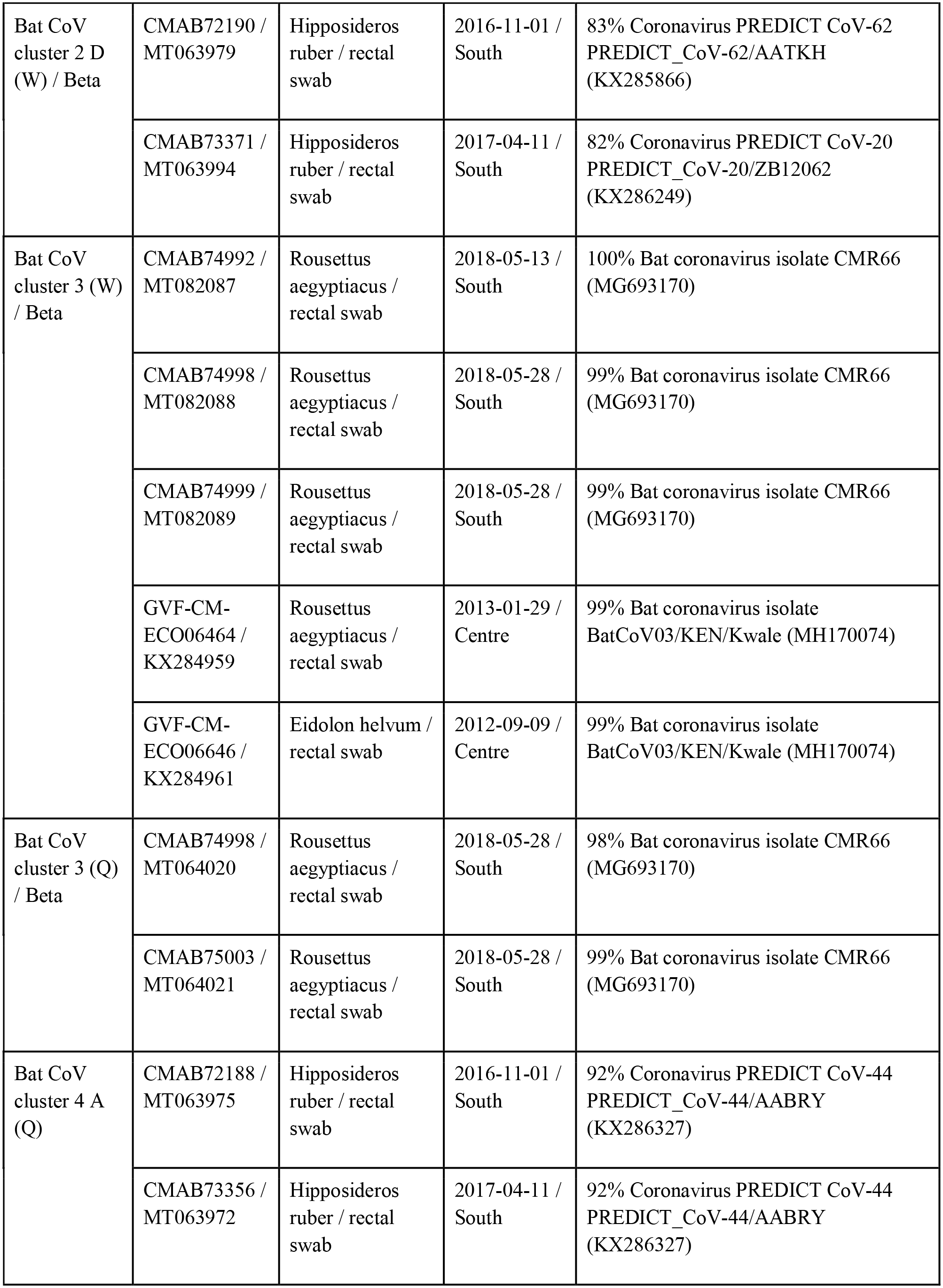
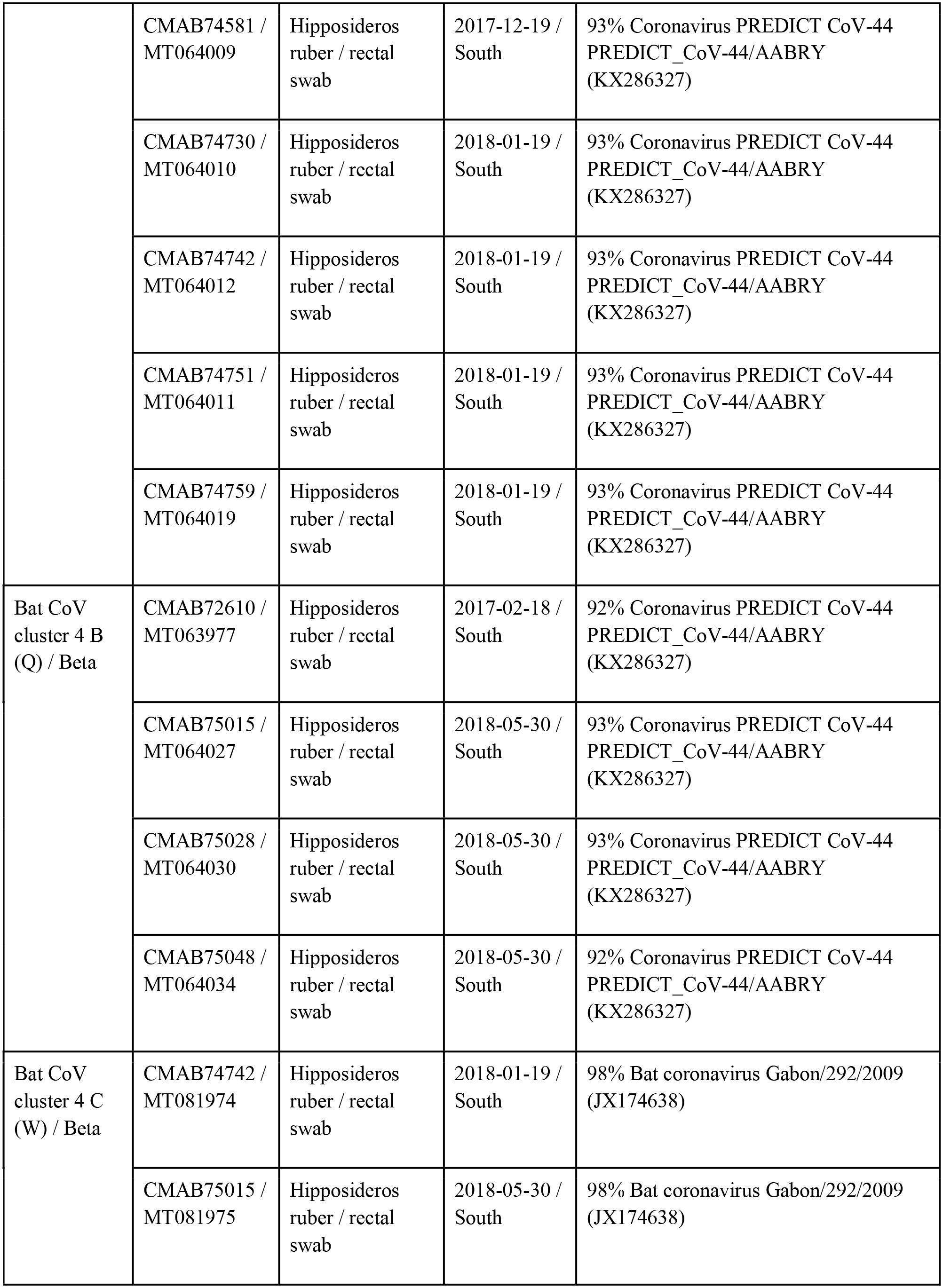
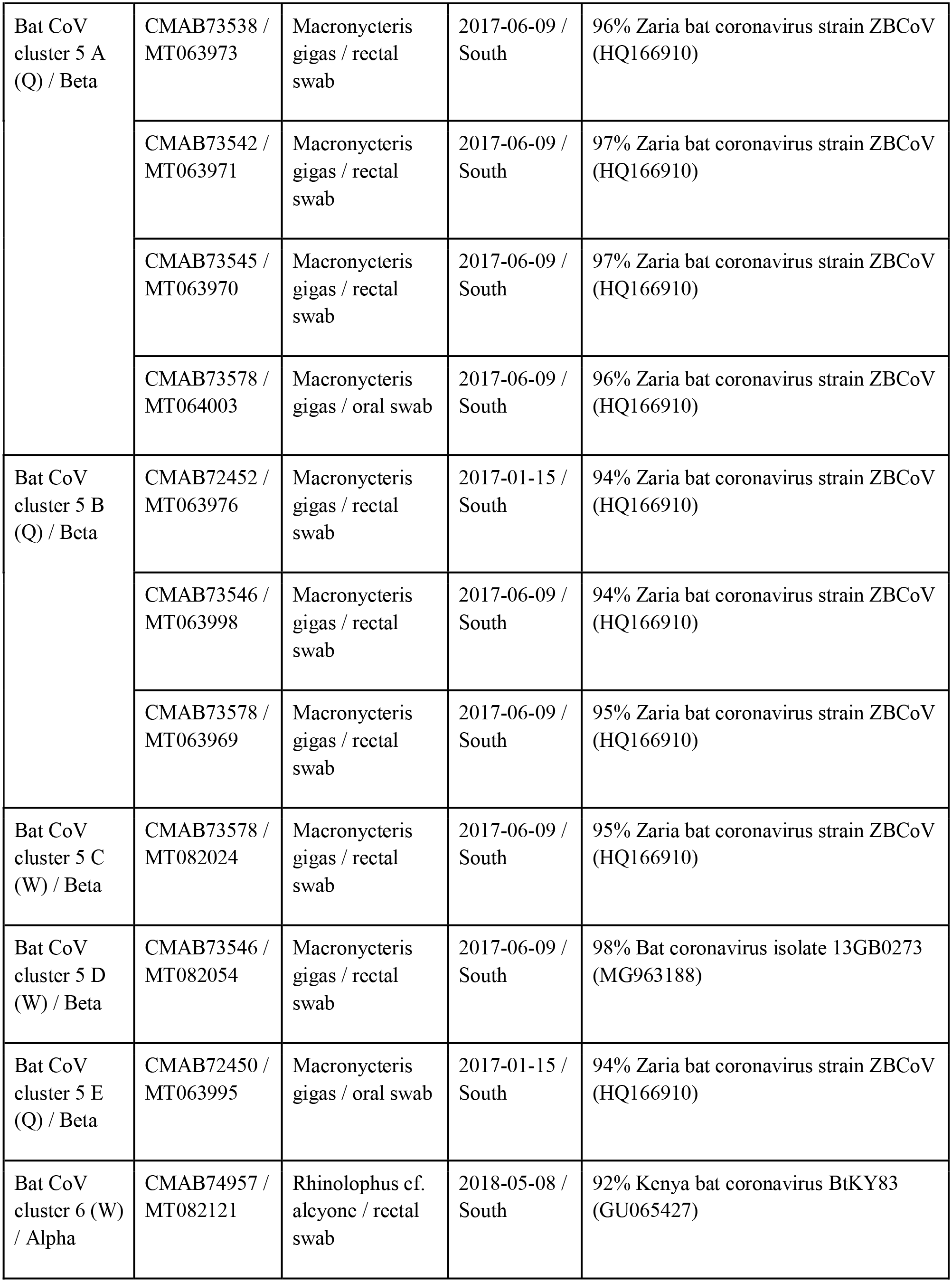
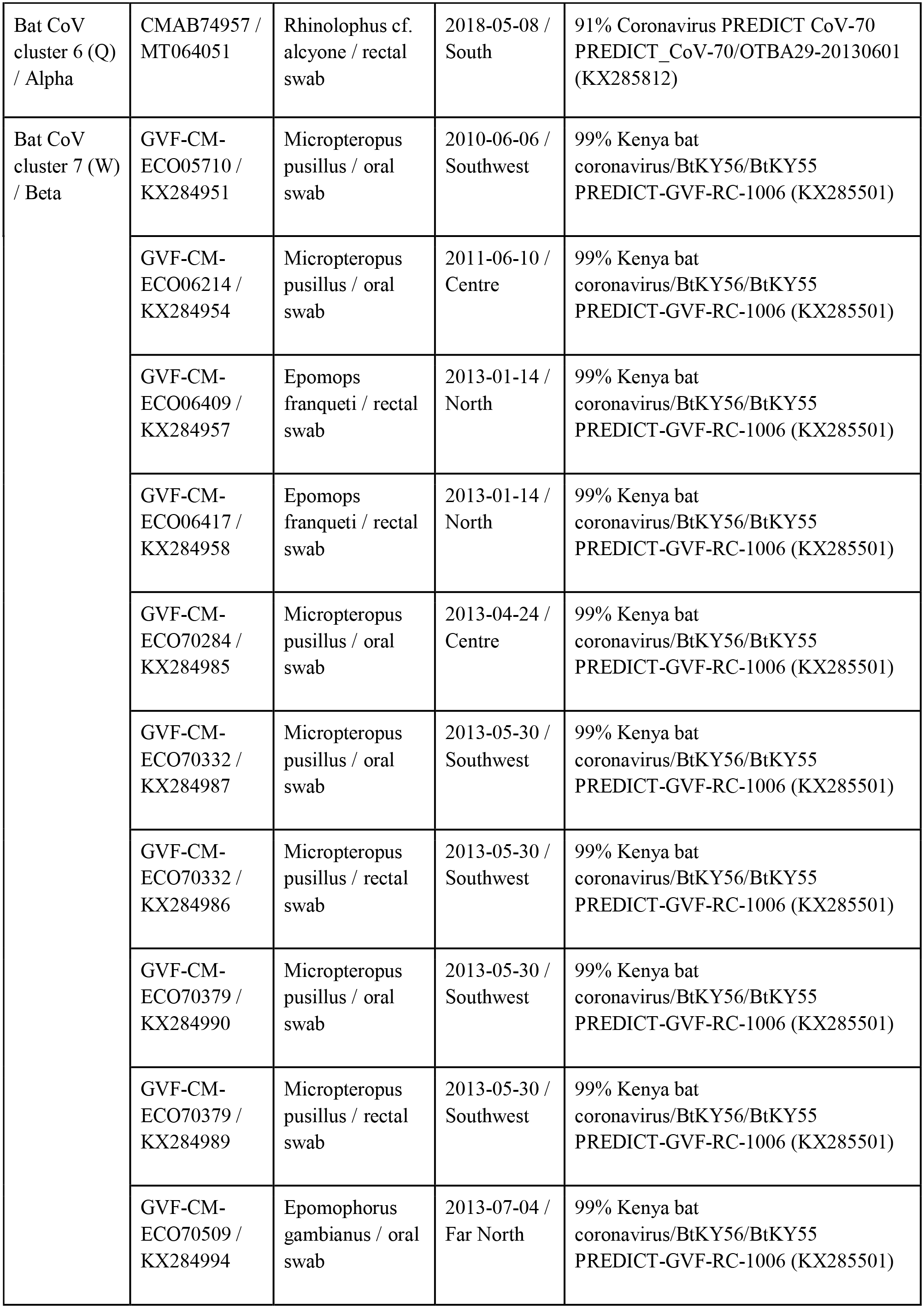
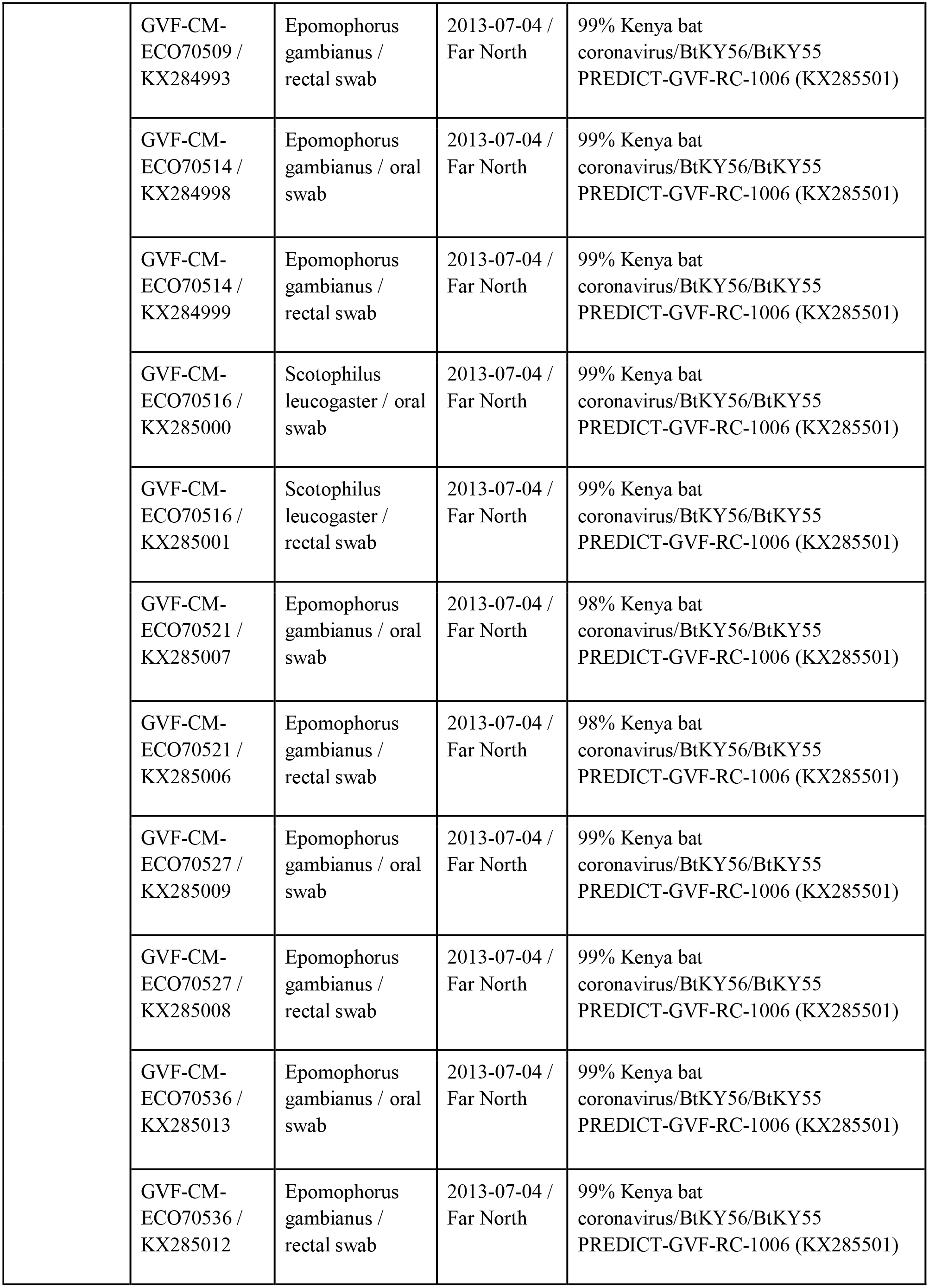
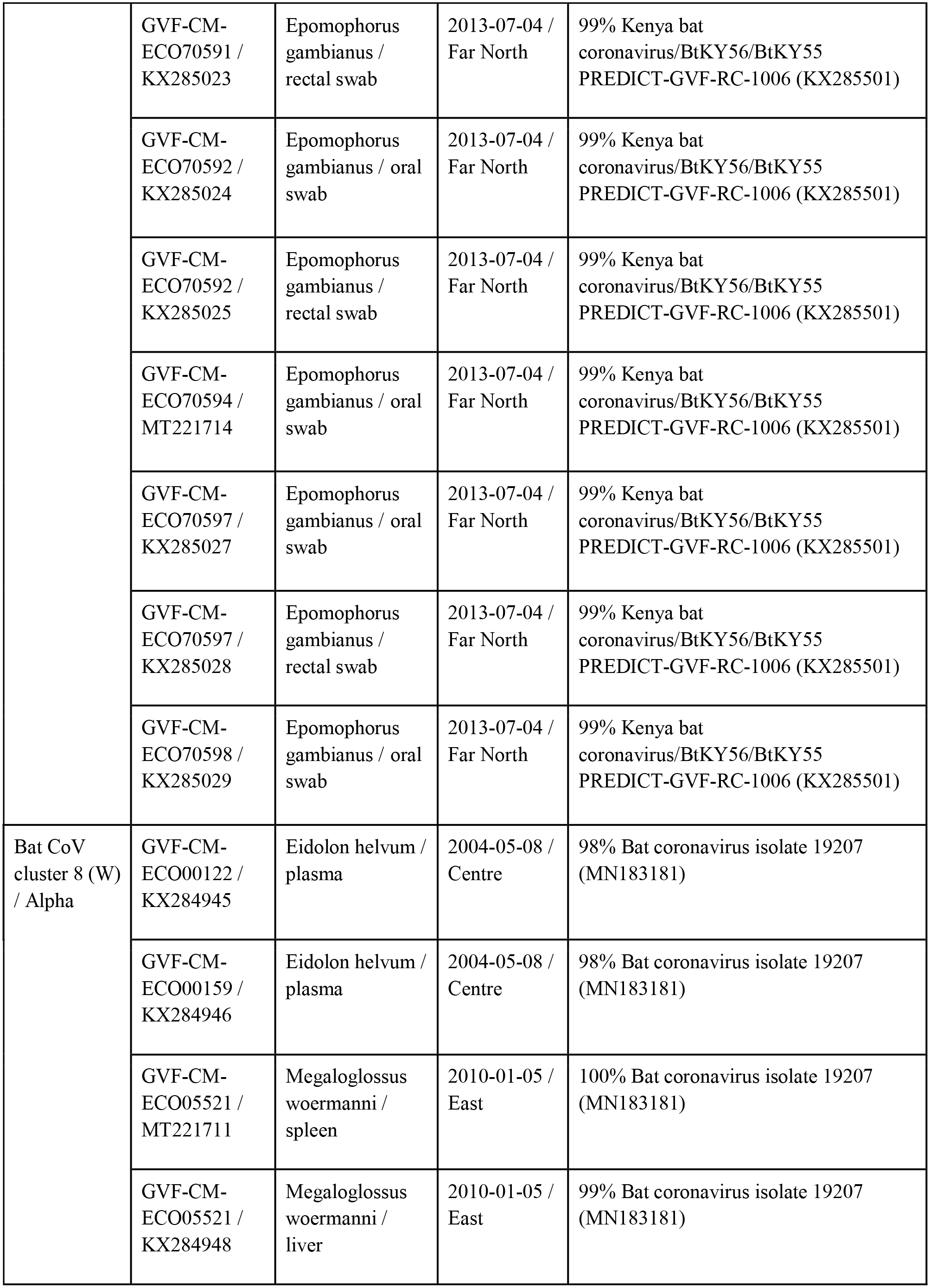
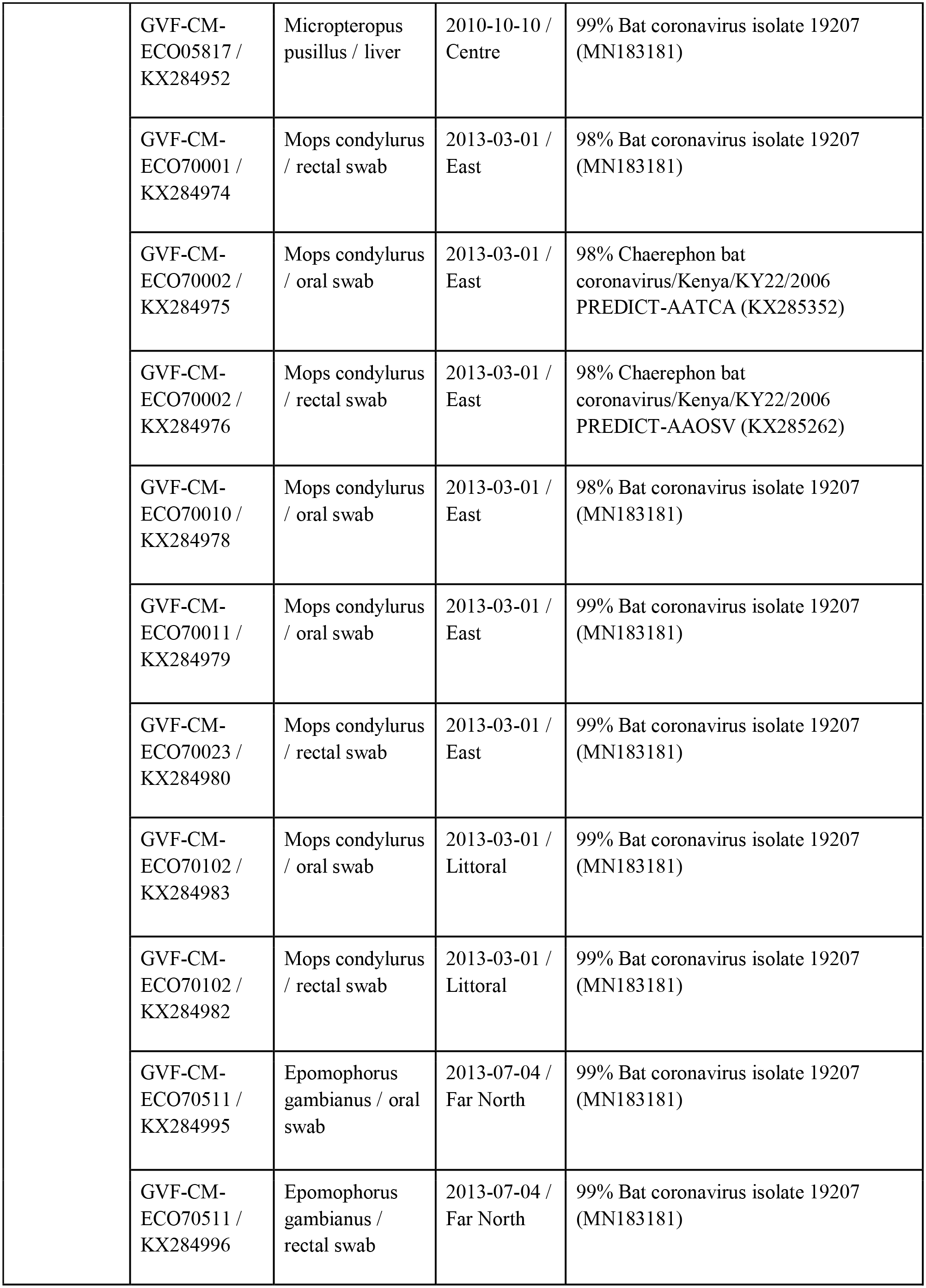
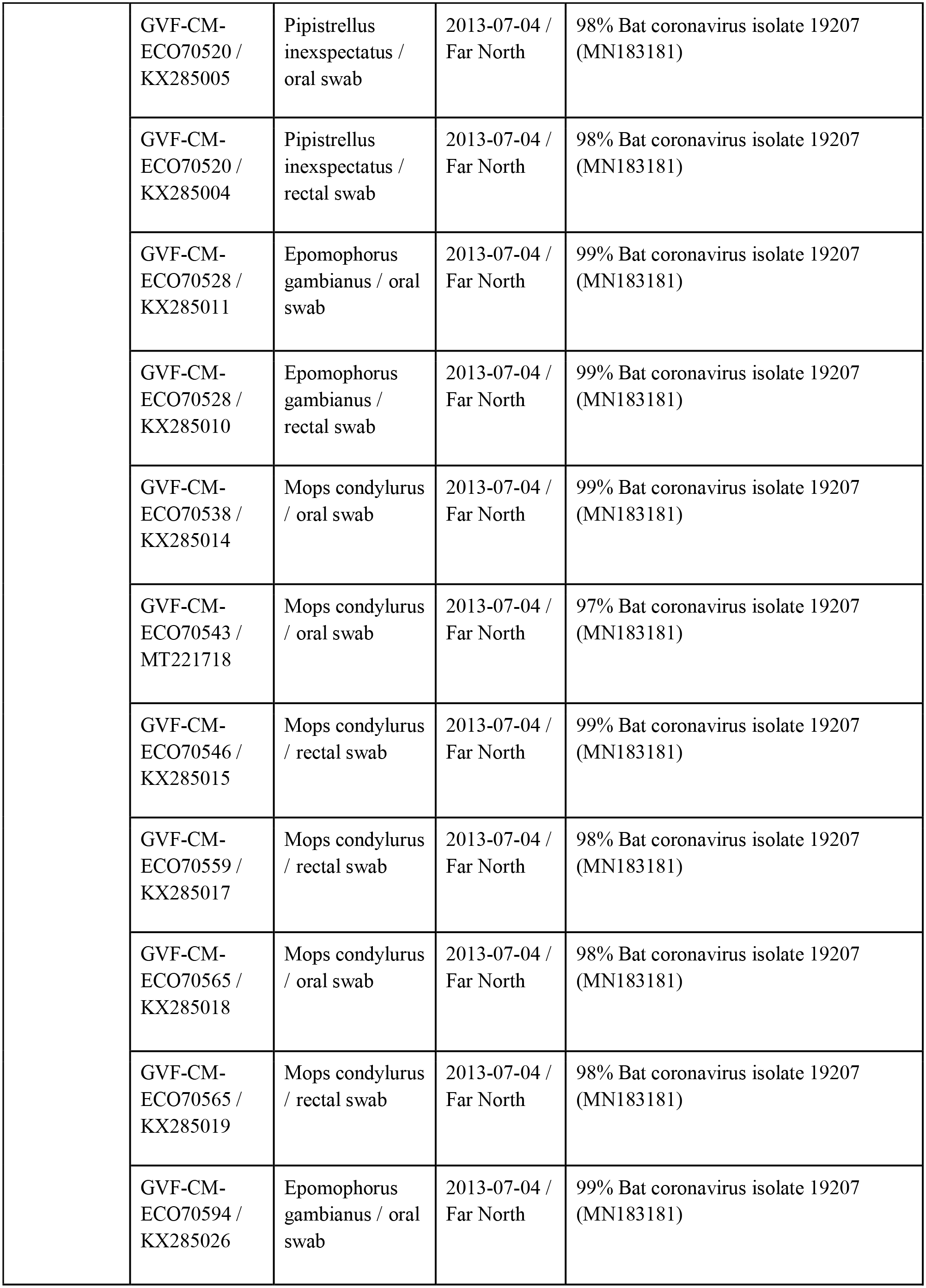
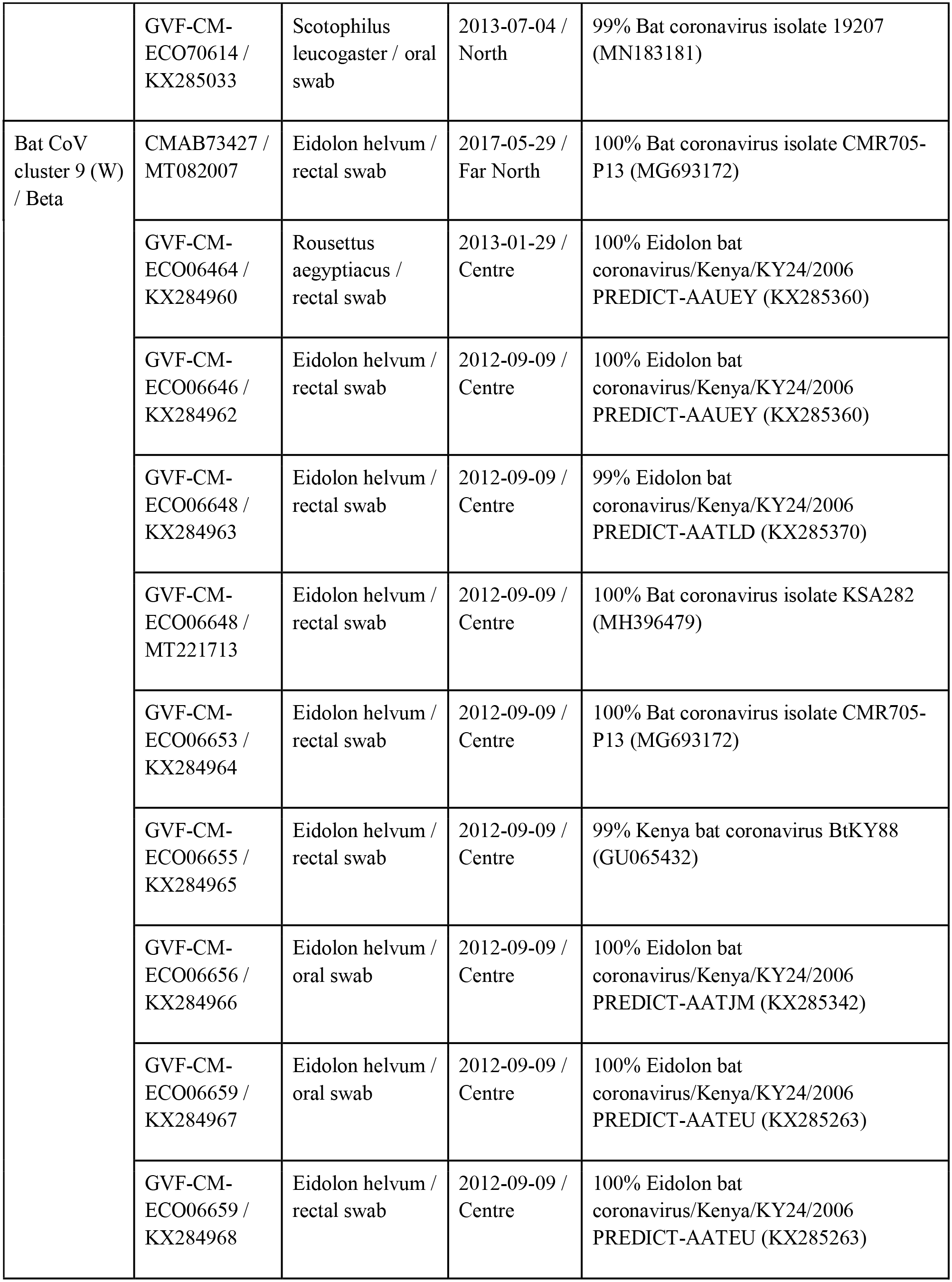
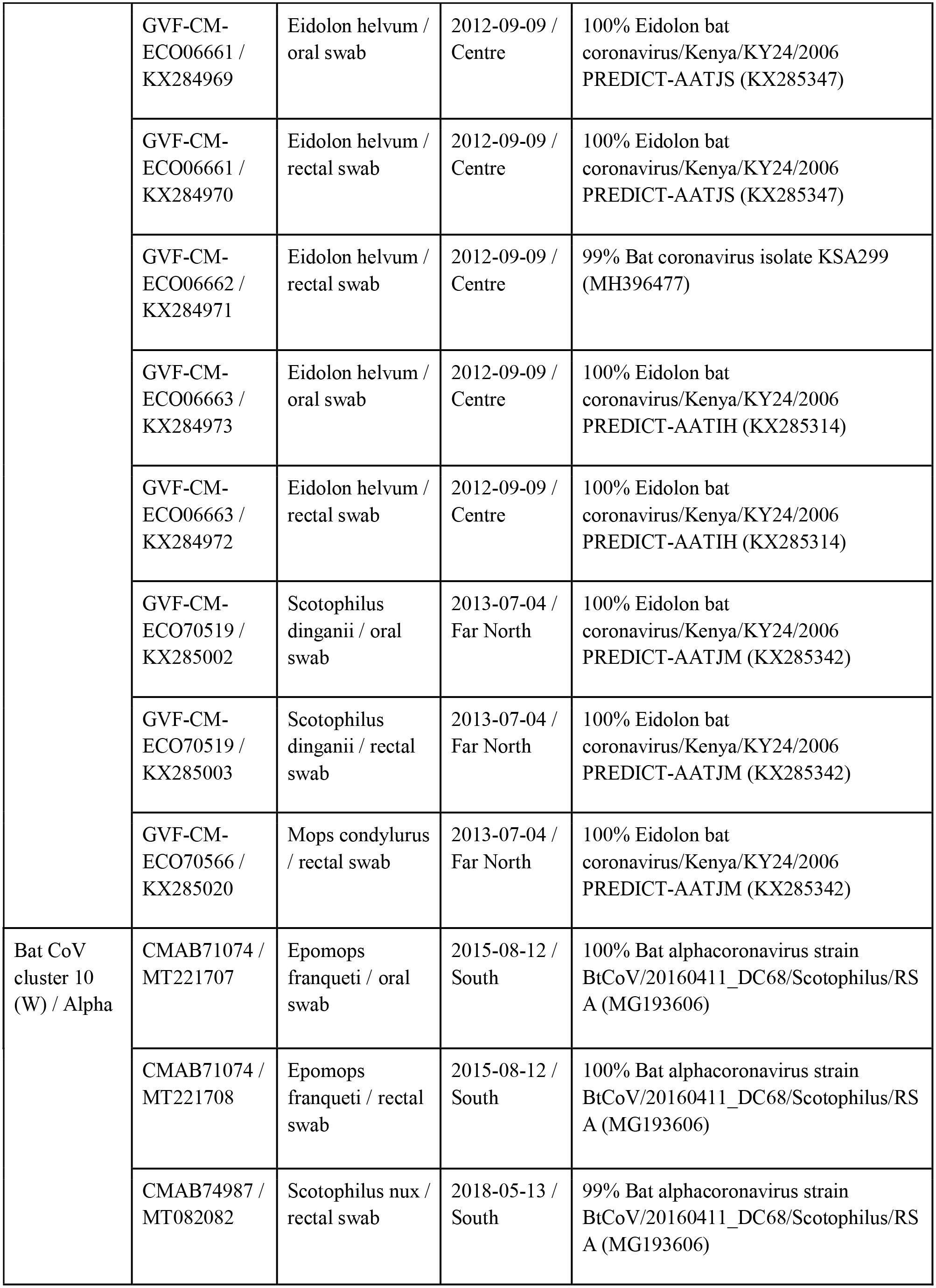
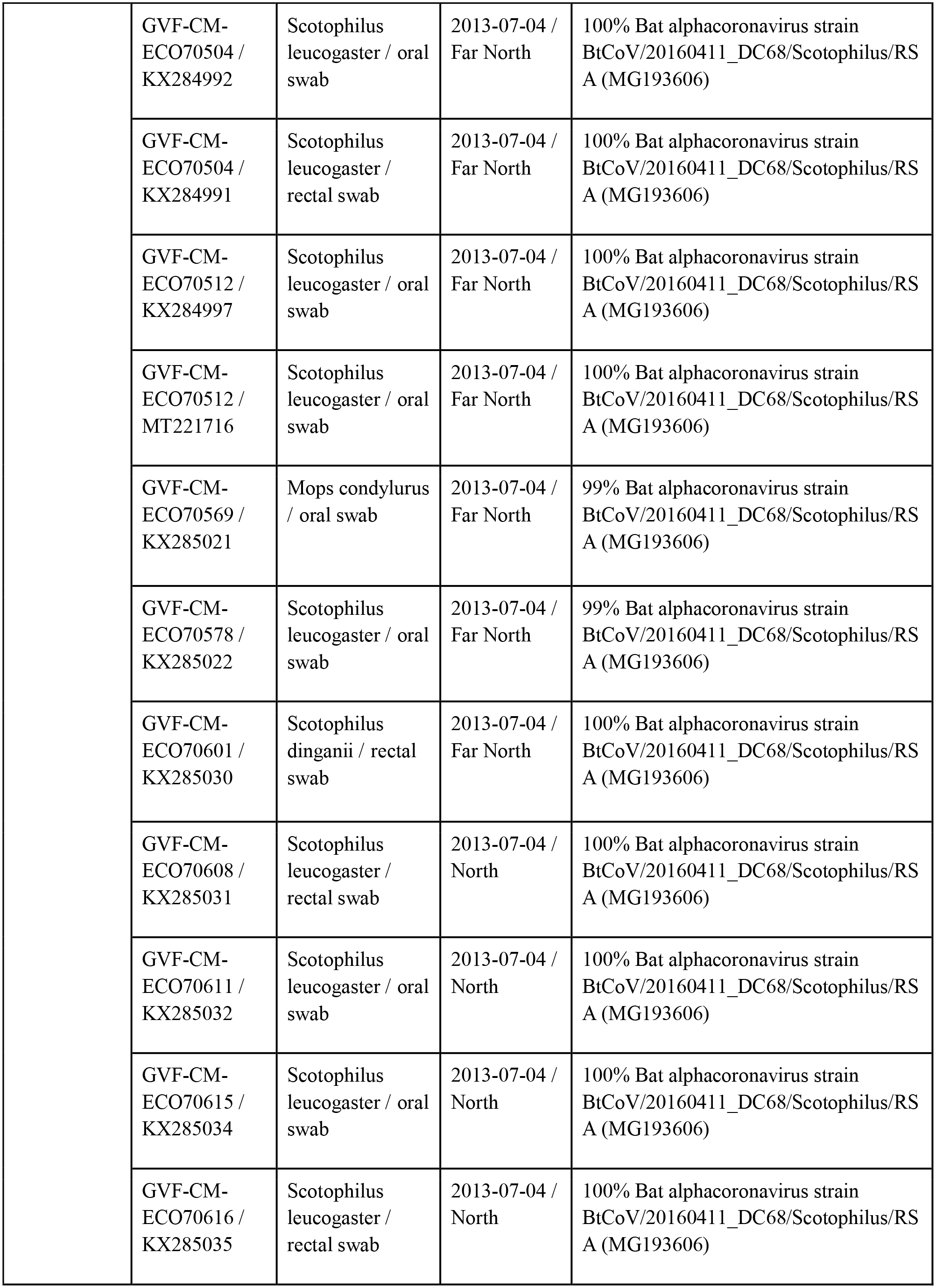
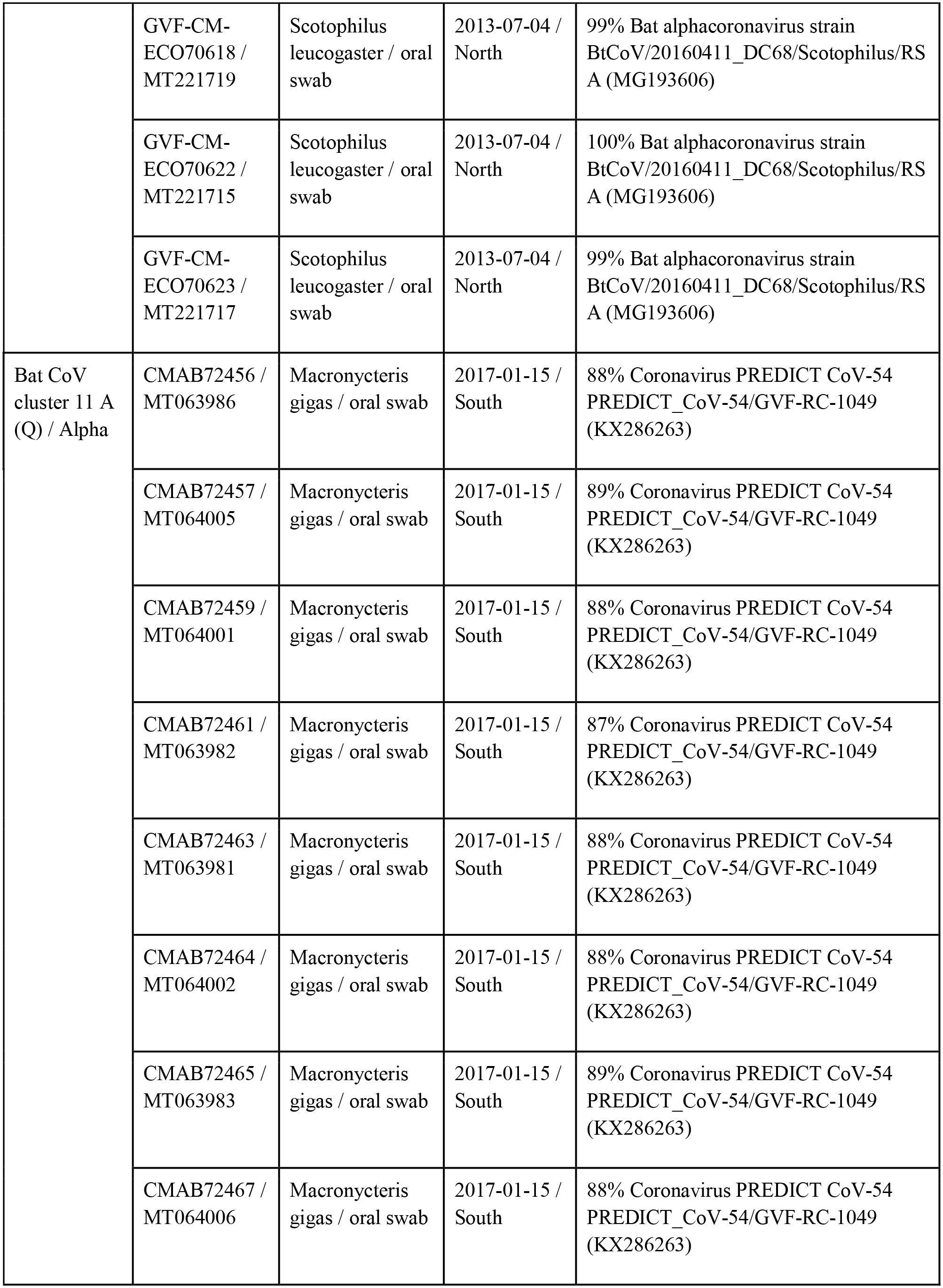
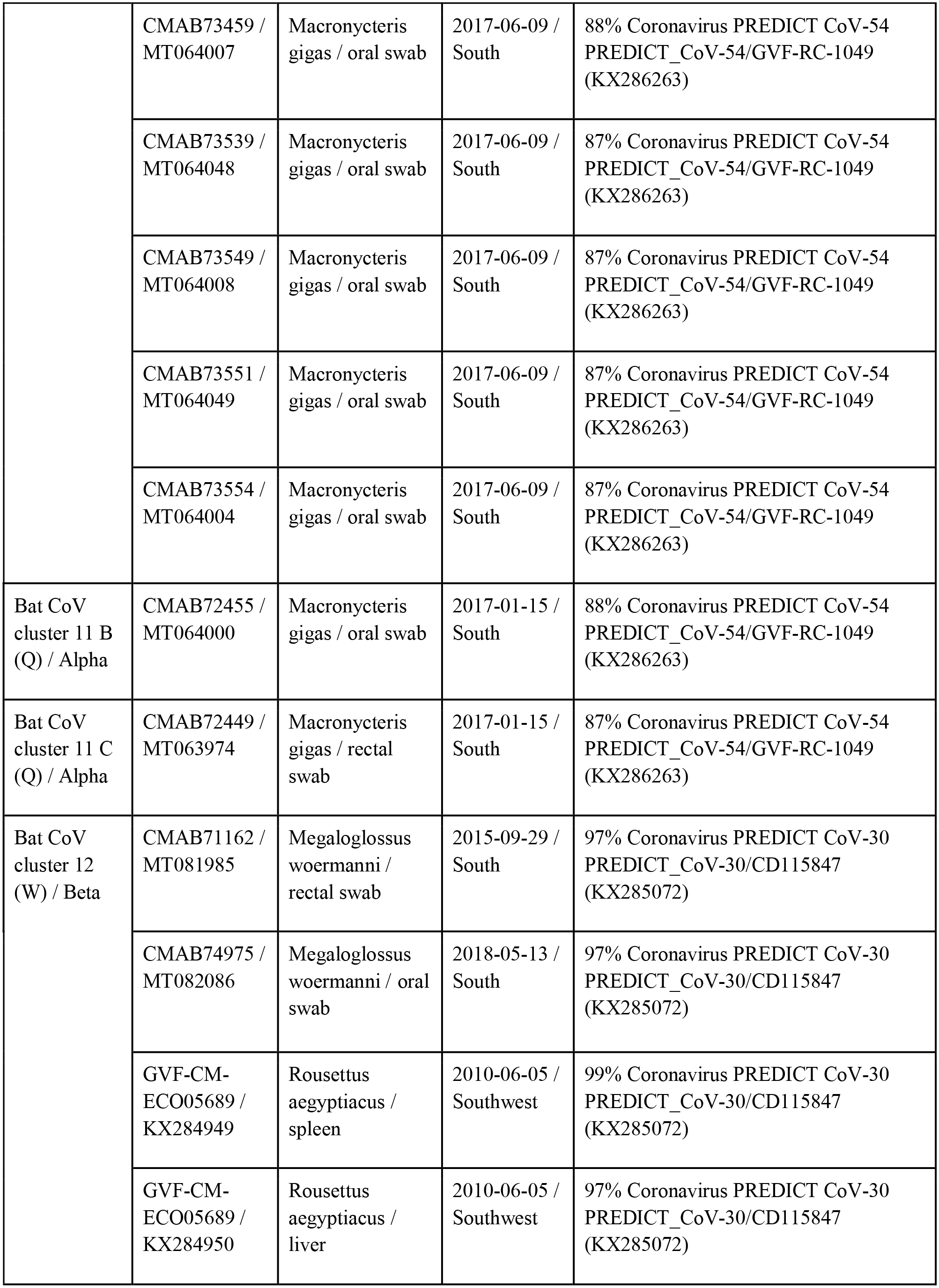
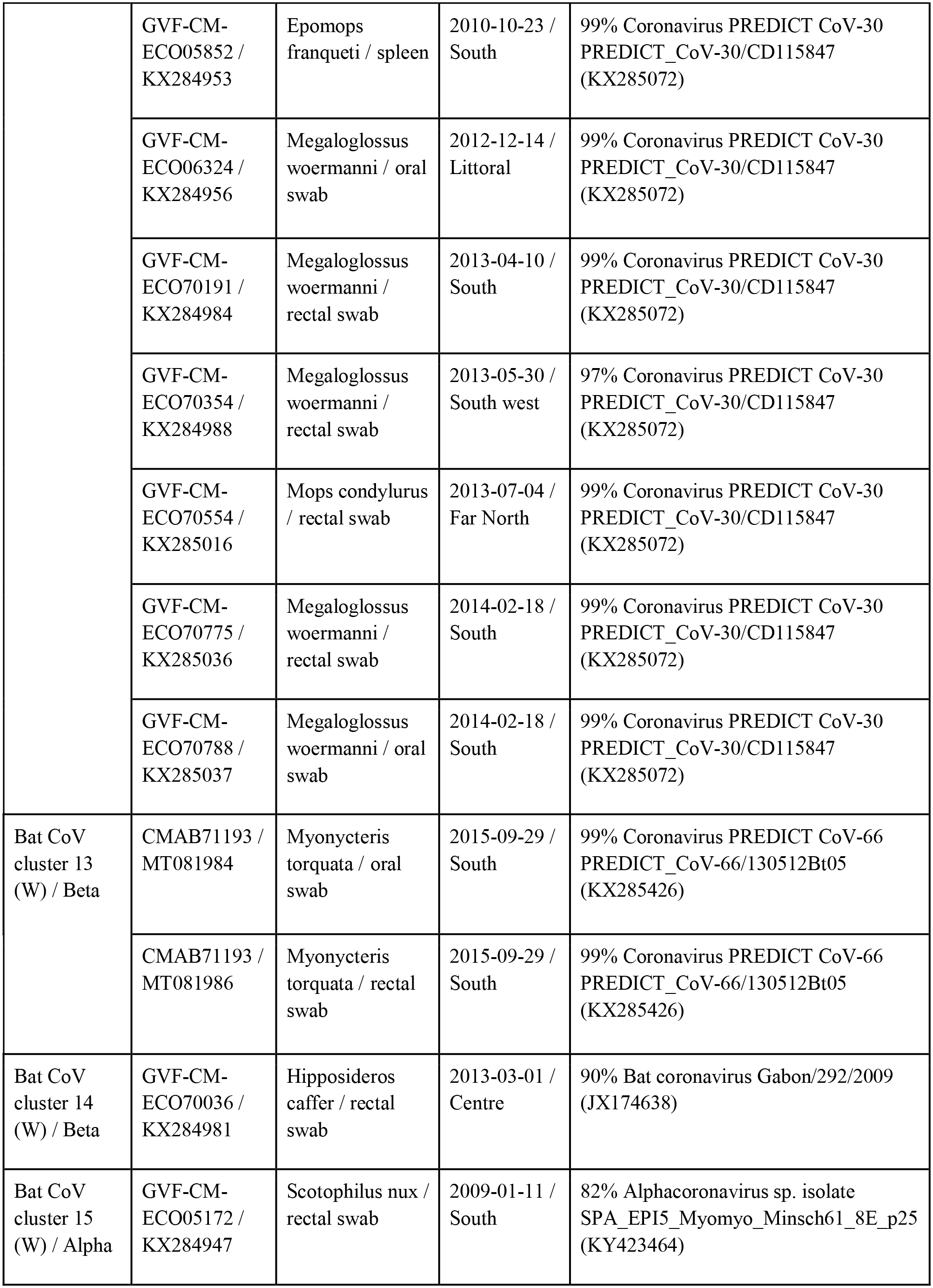
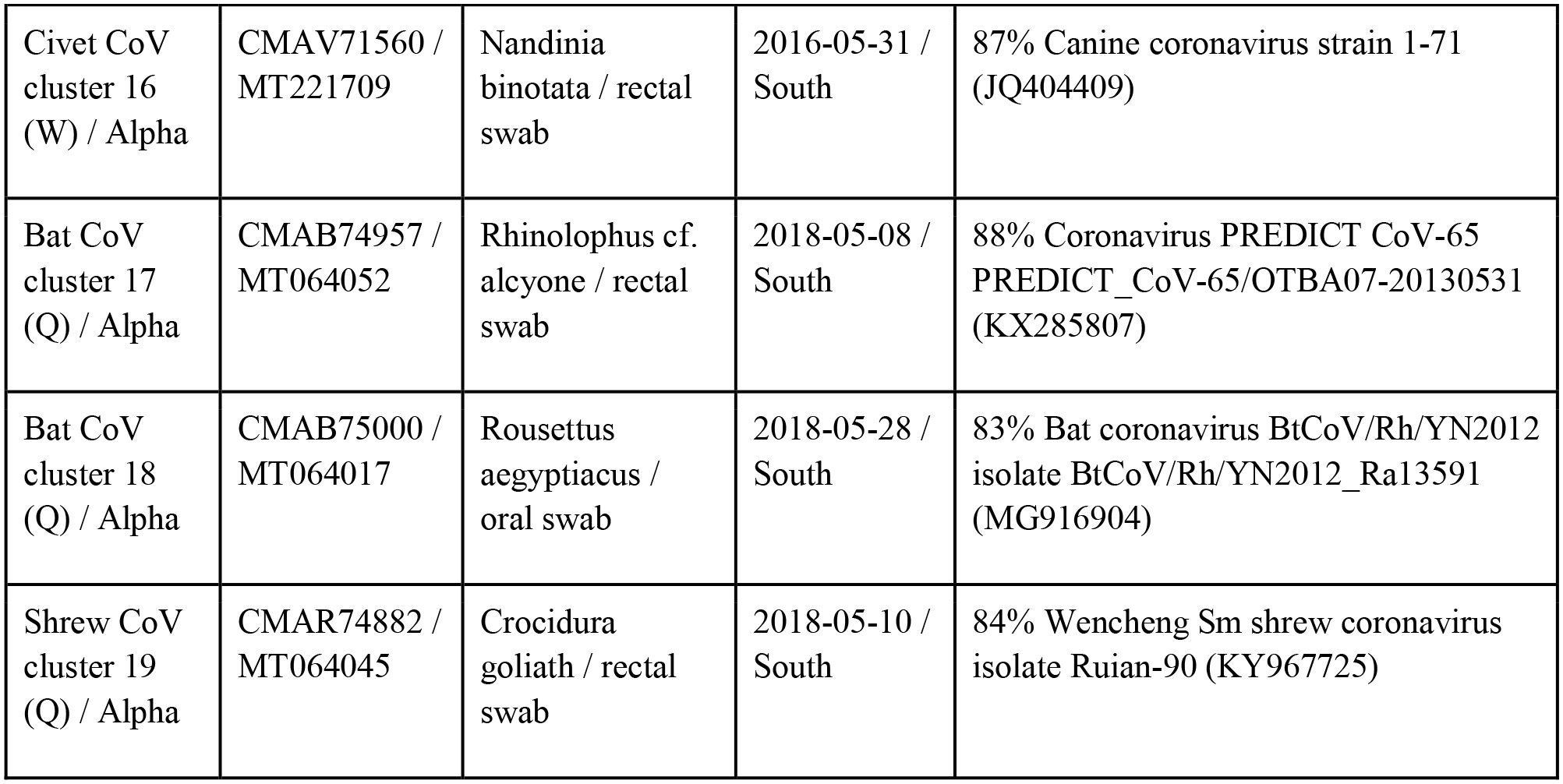
List of samples containing CoV RNA.

### Phylogenetic analysis

To facilitate phylogenetic analysis, select sequences of published complete CoV genomes isolated from humans, bats and other hosts, as well as partial sequences from CoVs closely related to those detected in this study were included. Sequences from novel Cameroonian isolates were included if they differed from others by at least 5%. Multiple sequence alignments were made in Geneious (version 11.1.3, ClustalW Alignment). Bayesian phylogeny of the polymerase gene fragment was inferred using MrBayes (version 3.2) with the following parameters: Datatype=DNA, Nucmodel=4by4, Nst=1, Coavion=No, # States=4, Rates=Equal, 2 runs, 4 chains of 10,000,000 generations. The sequence of an avian Gamma Coronavirus (NC_001451) served as outgroup to root the trees based on the RdRp PCR amplicons, while HCoV-NL63 (AY467487) served as outgroup for sequences related to HCoV-229E. Trees were sampled after every 1,000 steps during the process to monitor phylogenetic convergence, and the final average standard deviation of split frequencies was below the MrBayes recommended final average <0.01 for all analyses [Ronquist 2012]. The first 10% of the trees were discarded and the remaining ones combined using TreeAnnotator (version 2.5.1; http://beast.bio.ed.ac.uk) and displayed with FIGTREE (1.4.4; http://tree.bio.ed.ac.uk/) [Bouckaert 2019].

### Statistical analysis

Ecological data collected along with the samples obtained from bats was statistically analyzed in relation to the outcome of PCR tests. The variables included species, family, and suborder, age and sex, and factors such as the interface of exposure with humans, and season (wet/dry) of sampling. The taxonomy information for the analysis was based in the Integrated Taxonomic Information System (ITIS) website (https://www.itis.gov/), age coded as either adult or non-adult, interface categorized as ‘value chain’ for animal samples obtained at markets or directly from hunters, ‘tourism’ for samples obtained at zoos and sanctuaries, and ‘other peridomestic’. The seasons were defined as switching from wet to dry after November 15^th^ and from dry to wet after March 15^th^ of each year. All statistical data analyses were conducted using the statistical software SPSS (Statistical Package for the Social Sciences) version 26. At the univariate level, frequencies and percentages of the selected variables of interest (i.e. PCR test results, species, family, suborder, age, sex, interface, season, etc.) were generated. At the bivariate level, simple cross-tabulation, chi-square tests have been used to determine the statistical association (which are considered statistically significant at 5% level) between outcome and predictor variables. Here, the PCR test result has been considered as the dependent variable, and species, family, suborder, age, sex, interface, and season have been considered as the predictor/independent variables.

## Results

### Sample set

A total of 11,474 samples from 6,580 animals of several different orders were collected between 2003 and 2018, covering all 10 regions of Cameroon (Figure 1). Animals sampled included 2,740 rodents (28 species), 2,581 bats (50 species), 1,006 primates (24 species), 159 Eulipotyphla (3 species), 38 pangolins (1 species), 37 carnivores (3 species), 17 even-toed ungulates (4 species), and 2 hyraxes (1 species) (Supplement 2). Samples were predominantly oral (5,214) and rectal (4,818) swabs, but also included tissue such as spleen (893), liver (93), lung (5), colon (4), small intestine (4), muscle (1), skin (1) and thyroid (1). Other sample types included plasma (377), serum (20), and whole blood (1), as well as feces (36), genital swabs (5), and nasal swabs (1).

**Figure 1.**
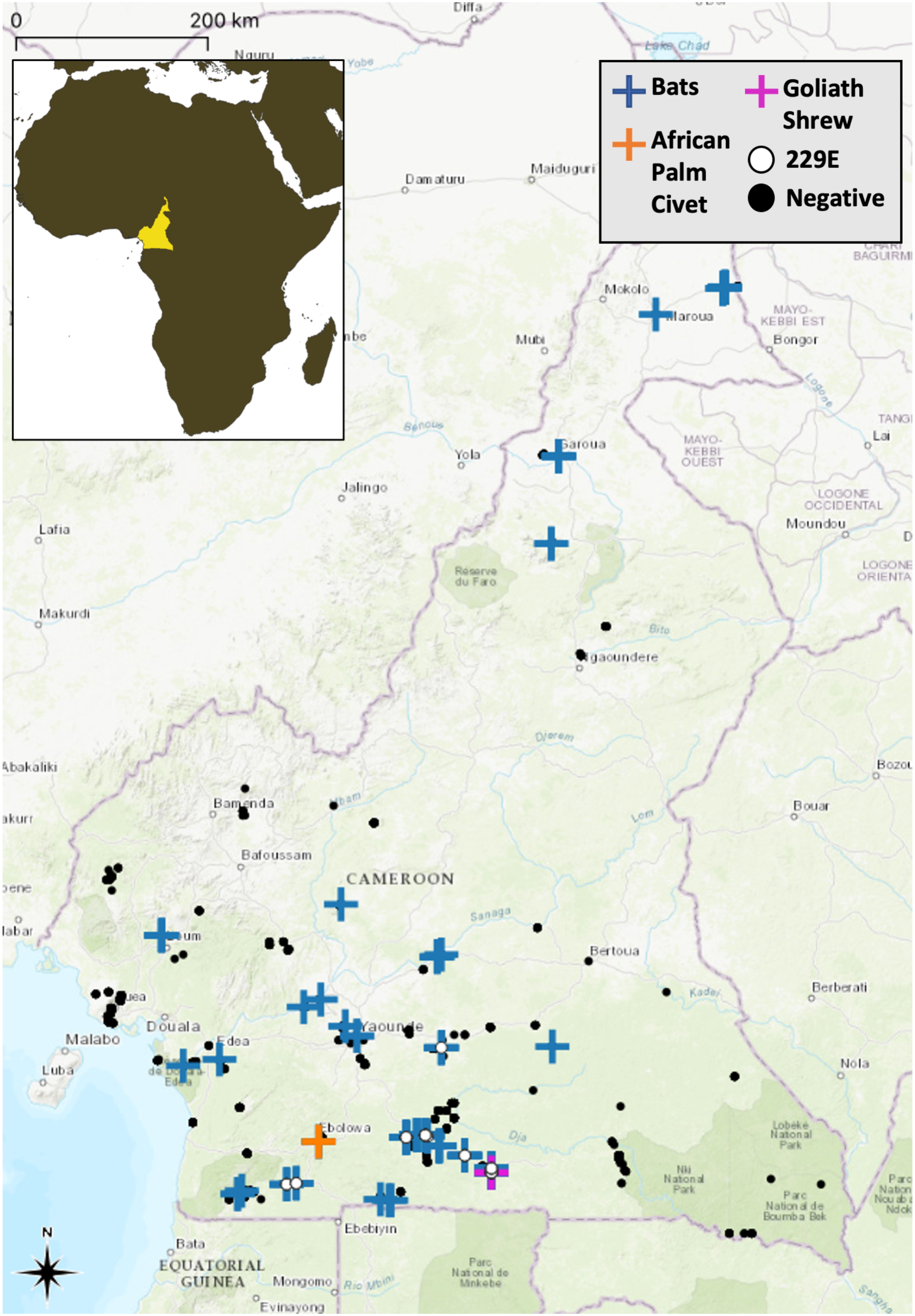
Sampling map: Map of Cameroon highlighting where samples were collected.

Coronavirus RNA was detected in at least one sample with at least one PCR assay in 175 individual bats, one civet, and one shrew (Table 2). Rectal swabs were CoV RNA positive in 129 instances, oral swabs in 71 instances, liver and spleen in 3 instances each and plasma in 2 instances. The Watanabe PCR protocol produced 173 positive results, while the Quan PCR protocols produced 64 positive results.

Bat sampling was conducted in many areas of the country, but 46.7% of the bats were sampled in the South Region (Supplement 3). Male bats were slightly overrepresented (53.5%) compared to females, however female bats were significantly (p<0.01) more likely to have a positive CoV test (8.2%) than male bats (5.6%). Differences in CoV RNA positive rate were also observed between adult (7.4%, n=1495) and younger bats (10.3%, n=87), but these were not statistically significant. Samples collected at animal-human interfaces categorized as ‘Tourism’ (1.5%, n=136) and ‘Value chain’ (3.1%, n=488) were significantly less likely (p<0.001) to be CoV RNA positive than those collected at ‘other peridomestic’ interfaces (8.1%, n=1957).

### Civet CoV

The CoV RNA detected in an African palm civet *(Nandinia binotata)* resembles that of an alphacoronavirus and, on the nucleotide level (BLASTN), is most closely related to canine CoV, feline CoV, and porcine CoV (TGEV) with 87%, 86%, and 85% identities respectively (Table 2). Phylogenetic analysis places the RNA in a cluster with dog, cat, mink, and pig CoVs (Figure 2).

**Figure 2.**
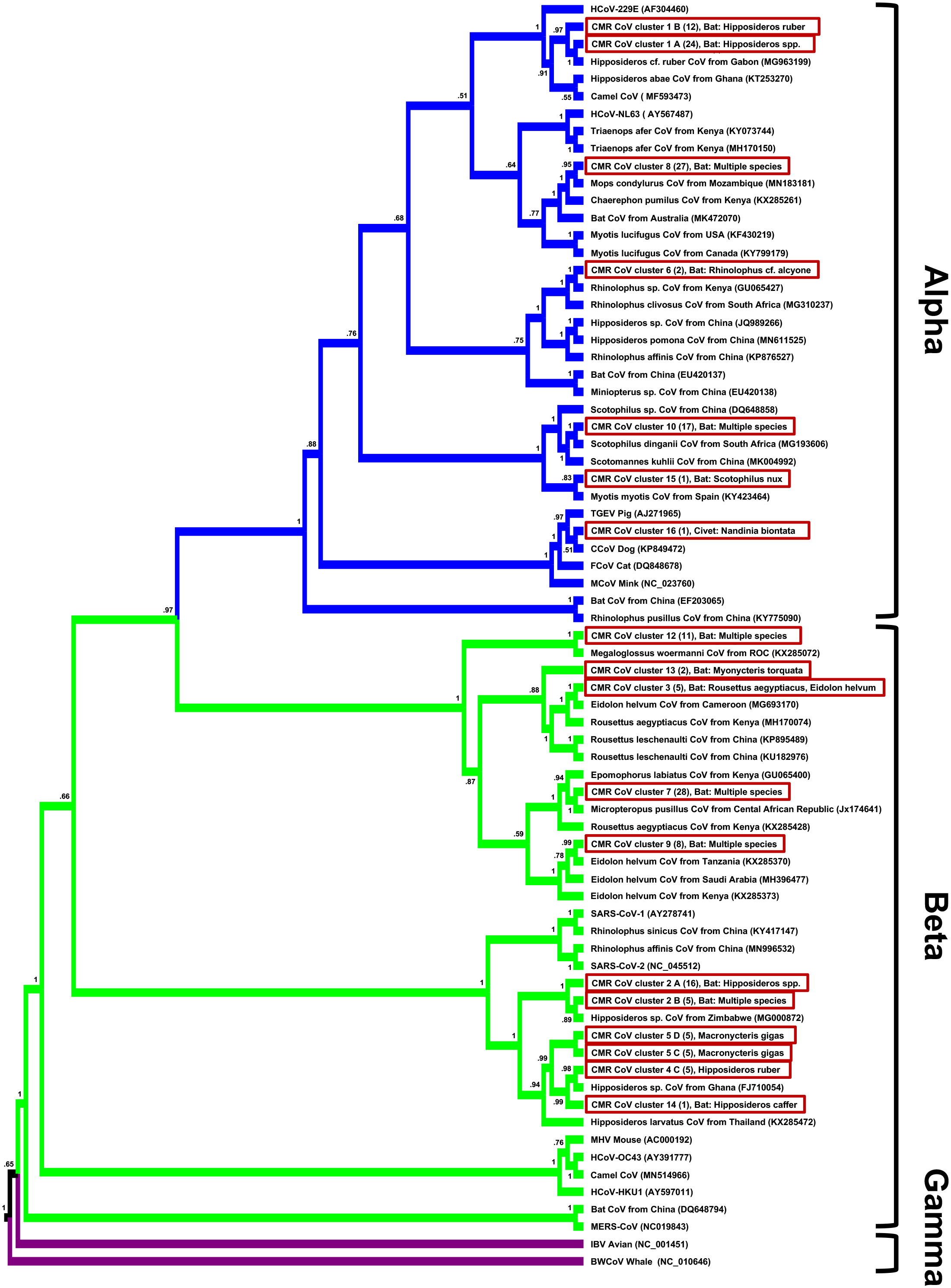
Phylogenetic tree: Maximum likelihood phylogenetic tree of coronavirus sequences presented as a proportional cladogram, based on the RdRp region targeted by the PCR by Watanabe et. al. [Watanabe 2010]. The sequences detected during the project are highlighted by red boxes and numbers in brackets indicate the number of sequences sharing more than 95% nucleotide identities. GenBank accession numbers are listed for previously published sequences, while sequences obtained during the project are identified by cluster names (compare Table 2). Numbers at nodes indicate bootstrap support.

### Shrew CoV

The CoV RNA detected in a Goliath shrew (*Crocidura goliath*) resembles that of an alphacoronavirus and is on the nucleotide level (BLASTN) closest related to isolates of Wencheng Sm shrew CoV (from a *Suncus murinus*), and isolates of Coronavirus PREDICT CoV-46 (from a *Crocidura* sp.), with up to 84% and 83% identities, respectively (Table 2). Phylogenetic analysis places the RNA in a cluster with other shrew CoVs as part of a basal branch of alphacoronaviruses (Figure 3).

**Figure 3.**
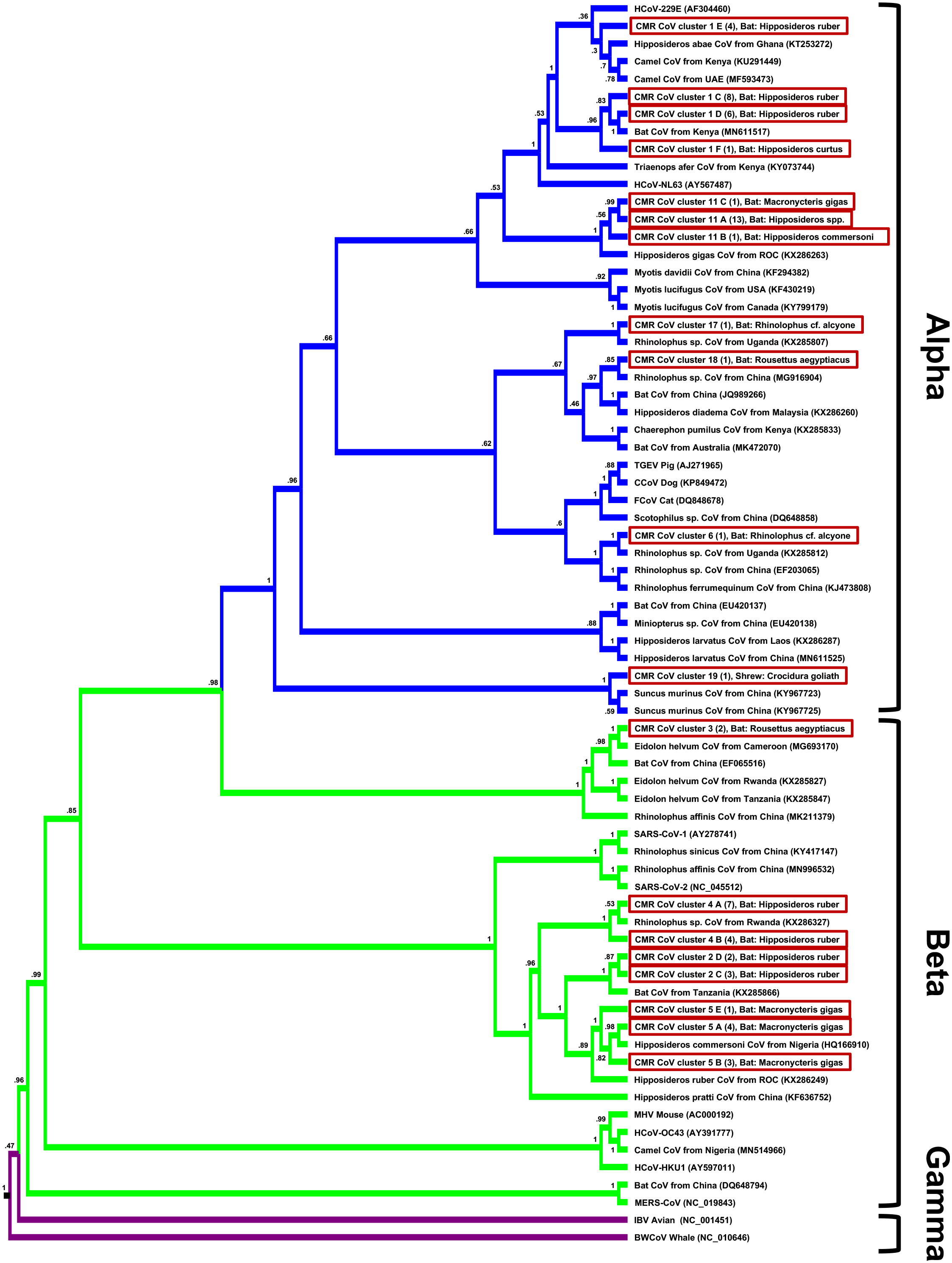
Phylogenetic tree: Maximum likelihood phylogenetic tree of coronavirus sequences presented as a proportional cladogram, based on the RdRp region targeted by the PCR by Quan et. al. [Quan 2010]. The sequences detected during the project are highlighted by red boxes and numbers in brackets indicate the number of sequences sharing more than 95% nucleotide identities. GenBank accession numbers are listed for previously published sequences, while sequences obtained during the project are identified by cluster names (compare Table 2). Red boxes indicate isolates from this study. Numbers at nodes indicate bootstrap support.

### Bat CoVs

The CoV RNA detected in the 175 bats form 17 different genetic clusters, of which 8 coincide with alpha and 9 with beta CoVs. The majority of sequences share identities above 90% with known bat CoVs, while the sequences in up to 6 of the 17 clusters do not (bat CoV clusters 2, 11, 14, 15, 17, 18) (Table 2, Figures 2 and 3). RNA corresponding to a single CoV was detected in 170 bats. In 89 bats the CoV RNA resembled that of an alpha CoVs, and in 81 cases that of a beta CoV. In 5 bats RNA corresponding to two different CoVs was found. In two of these cases, RNA of both an alpha and a beta CoV was present, in two bats the RNA of two different beta CoVs was detected, and in one bat we found the RNA of two different alpha CoVs (Table 2). In all other cases (n=30) where we detected RNA in more than one sample of the same bat, the RNA amplified by the same PCR differed by less than 10%. In 24 instances they were 100% identical, in five instances they differed by less than 1% and in one instance the difference was 5.2%.

### HCoV-229E-like sequences in bats

Sequences resembling HCoV-229E were found in 40 bats from the *Hipposideridae* family, primarily in the species *Hipposideros ruber*, and showed differences of up to 6% among each other in the Watanabe amplicons and up to 11% in the Quan amplicons. Additional sequence information of the full or partial S, E, M, and N genes was obtained, and differences in the sequences ranged up to 8% for the E, 12% for the M, 32% for the N and 26% for the S genes. Phylogenetic clustering of sequences differed depending on the gene, with isolates BtCoV/KW2E-F56/Hip_cf._rub/GHA/2011 (KT253271) and BtCoV/AT1A-F1/Hip_aba/GHA/2010 (KT253272) being consistently the most basal isolates in the CoV-229E branch (Figure 4, Supplement 1). The phylogenetic analyses, that involve sequence isolates from bat, camel, and human hosts, place the HCoV-229E isolates that were obtained from humans closest to isolates from camels in case of the S and N genes, but closest to isolates from bats for the E, M and RdRp genes (Figure 4, Supplement 4).

**Figure 4.**
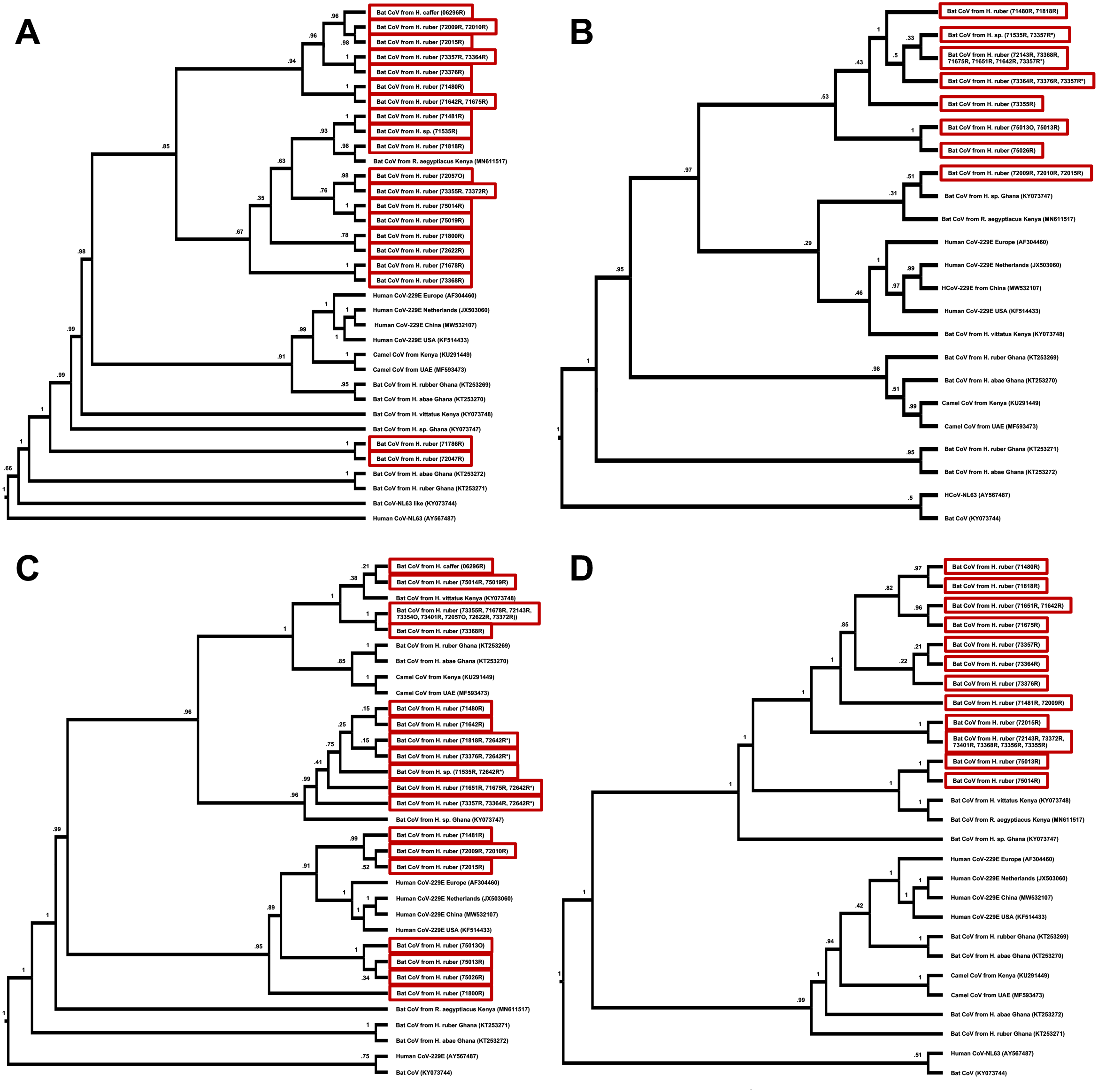
Phylogenetic trees of HCoV-229E-like isolates: Maximum likelihood phylogenetic tree of coronavirus sequences related to HCoC-229E, based on the Spike (A), Envelope (B), Membrane (C) and Nucleoprotein (D). Red boxes indicate isolates from this study. Numbers at nodes indicate bootstrap support. Compare also Supplement 4.

### Seasonality and other predictors in bats

Bat sample collection was focused on seasons, and thus varied over the months, with a high of 432 samples collected in March and a low of 48 in August (Supplement 5). Bats sampled in the wet season accounted for 61.9% of total bats compared to 38.1% in the dry season. The proportion of CoV RNA positive bats varied significantly (p=<0.001) between the wet (8.2%) and dry season (4.5%), while the proportion of events (samples collected at the same location on the same day) with at least one positive animal was very similar with 29.4% during the wet season and at 28.3% during the dry season. The proportion of CoV RNA positive individuals for the sampling events with >25 bats (n=24) fluctuated between 0% and 48%, and was at 21.2% for the largest event. Nine events took place during the wet season, and 15 during the dry season. Among the bats sampled during this largest event were 17 different species with varying rates of CoV RNA detection, including *Chaerephon pumilus* (0%, n=29), *Epomophorus gambianus* (54.5%, n=22), *Mops condylurus* (26.7%, n=30), *Nycteris hispida* (0%, n=18), *Scotophilus dinganii* (16.7%, n=12), and *Scotophilus leucogaster* (38.7%, n=31).

Seasonal differences in the CoV RNA detection rate were associated with and differed depending on certain taxonomic species, families, and suborders (Table 3). Both *Yinpterochiroptera* and *Yangochiroptera* bats were overall more likely to be CoV RNA positive during the wet season, with the observed difference being more pronounced in the latter. Significant seasonal differences were observed in five species; high rates of CoV RNA detections were associated with the wet season and low rates with the dry season for *Eidolon helvum*, *Hipposideros ruber*, and Mops condylurus, while it was the opposite for *Macronycteris gigas* (Table 3).

**Table 3:** PCR results of suborder, family, and species by season (bats)

| *Suborder, family (≥10 sampled individuals) and species (≥10 sampled individuals)* | **Wet Season**<br>**PCR positives** | **Dry Season**<br>**PCR positives** | **Total**<br>**PCR positives** |
| --- | --- | --- | --- |
| ***Yinpterchiroptera total*\*\*\*** | **7.6% (107/1412)** | **4.5% (36/796)** | **6.5% (143/2208)** |
| ***Pteropodidae total*\*\*\*** | **6.1% (43/710)** | **1.6% (7/434)** | **4.4% (50/1144)** |
| *Eidolon helvum* ** | 10.6% (12/113) | 0.0% (0/154) | 4.5% (12/267) |
| *Epomophorus gambianus* | 37.5% (12/32) | - (0/0) | 37.5% (12/32) |
| *Epomops franqueti* | 1.5% (2/134) | 2.8% (2/72) | 1.9% (4/206) |
| *Lissonycteris angolensis* | 0.0% (0/13) | 0.0% (0/49) | 0.0% (0/62) |
| *Megaloglossus woermanni* | 3.6% (4/111) | 5.5% (4/73) | 4.3% (8/184) |
| *Micropteropus pusillus* | 5.1% (6/117) | 0.0% (0/20) | 4.4% (6/137) |
| *Myonycteris torquata* | 3.1% (1/32) | 0.0% (0/5) | 2.7% (1/37) |
| *Rousettus aegyptiacus* | 4.1% (6/147) | 1.9% (1/53) | 3.5% (7/200) |
| ***Hipposideridae total*** | **9.3% (63/681)** | **8.2% (29/354)** | **8.9% (92/1035)** |
| *Doryrhina cyclops* | 0.0% (0/34) | 0.0% (0/3) | 0.0% (0/37) |
| *Macronycteris gigas* *** | 8.3% (10/120) | 27.9% (12/43) | 13.5% (22/163) |
| *Hipposideros ruber* ** | 11.9 (50/421) | 6.5% (15/232) | 10.0% (65/653) |
| *Hipposideros caffer* | 1.1% (1/93) | 1.4% (1/72) | 1.2% (2/165) |
| ***Rhinolophidae total*** | **5.9% (1/17)** | **0.0% (0/8)** | **4.0% (1/25)** |
| *Rhinolophus landeri* | 0.0% (0/12) | 0.0% (0/7) | 0.0% (0/19) |
| ***Yangochiroptera total*\*\*\*** | **13.0% (24/184)** | **4.3% (8/188)** | **8.6% (32/372)** |
| ***Molossidae total*\*** | **11.4% (8/70)** | **4.8% (7/145)** | **7.0% (15/215)** |
| *Chaerephon pumilus* | 0.0% (0/31) | - (0/0) | 0.0% (0/31) |
| *Mops condylurus* *** | 22.2% (8/36) | 5.0% (7/139) | 8.6% (15/175) |
| ***Nycteridae total*** | **0.0% (0/30)** | **0.0% (0/17)** | **0.0% (0/47)** |
| *Nycteris grandis* | 0.0% (0/8) | 0.0% (0/15) | 0.0% (0/23) |
| *Nycteris hispida* | 0.0% (0/21) | 0.0% (0/2) | 0.0% (0/23) |
| ***Vespertilionidae total*\*** | **20.5% (16/78)** | **4.3% (1/23)** | **16.8% (17/101)** |
| *Neoromicia tenuipinnis* | 0.0% (0/6) | 0.0% (0/7) | 0.0% (0/13) |
| *Scotophilus dinganii* | 16.7% (2/12) | - (0/0) | 16.7% (2/12) |
| *Scotophilus leucogaster* | 38.7% (12/31) | 0.0% (0/2) | 36.4% (12/33) |
| **Total\*\*\*** | **8.2% (131/1597)** | **4.5% (44/984)** | **6.8% (175/2581)** |

\* Significant difference between calendric seasons $P < 0.10$ (Chi-square with Yates correction)
\*\* Significant difference between calendric seasons $P < 0.05$ (Chi-square with Yates correction)
\*\*\* Highly significant difference between calendric seasons $P < 0.01$ (Chi-square with Yates correction)

## Discussion

### Unexplored diversity

While we sampled and tested roughly equal proportions of bats and rodents and a significant number of NHPs in this study, all but 2 of the 177 animals that turned out to be positive for CoV RNA were bats. This is not a surprise, but it rather reinforces the notion that bats are the major reservoir for alpha- and beta-CoVs, and generally for viruses with zoonotic potential [Li 2005; Woo 2009; Annan 2013; O’Shea 2014; Anthony 2017]. While it has been suggested that under-sampling of rodents, rather than a particularly high prevalence in bats, may be a reason for this pattern, we did not find any evidence to support such a hypothesis in our study population, despite a high number of sampled rodents. However, considering the high diversity of rodent species, our sample is certainly not representative and biased towards those that thrive in a peridomestic environment or that are being hunted for consumption. The detection of novel CoV RNA in a civet is certainly an interesting finding, as civets played a key role as intermediate hosts in the emergence of SARS-CoV-1 [Kan 2005]. While civets are not farmed in Cameroon, they are hunted for consumption, which implies human contact, thus posing a potential risk. The detected RNA suggests a close relationship with other carnivore CoVs, and thus potentially a low risk for humans (Spillover Risk score of 54 out of 155), however in the absence of a full genomic sequence and further characterization experiments, this remains to be determined [Grange 2021].

The CoV RNA we detected belongs to 19 different genetic clusters, of which 8 might represent novel CoV species - six of these were found in bats, one in a civet, and one in a shrew. While we were unable to obtain large or complete genomic sequences from the isolates, these findings indicate that the CoV diversity in African wildlife species, and particularly in bats, is still poorly understood. This is concerning since spillover events are likely to occur, given the close interactions that arise from human housing conditions, hunting practices, ecotourism, and other human behaviors in Cameroon and other parts of Africa. While the emergence of SARS-CoV-1 and SARS-CoV-2 have put the spotlight on Southeast Asia and China in particular, it is important to keep in mind that the risks are present worldwide and local efforts should be enhanced across the globe to determine how to mitigate these risks [Jones 2008; Wolfe 2007]. CoV diversity poses a significant challenge for surveillance efforts and requires further exploration, but it also provides great opportunities to learn more about the biology and history of CoVs, including that of non-SARS CoVs, such as HCoV-229E.

### Viruses closely related to HCoV-229E are highly prevalent in Hipposideros bats

We detected RNA of bat CoVs closely related to HCoV-229E in 40 bats, which all belonged to the *Hipposideridae* family (3.9%, n=1,035). This finding, along with the fact that previous detections of HCoV-229E-like bat viruses were also almost exclusively associated with *Hipposideros* bats, supports the hypothesis that this family of bats constitutes the original host family and reservoir of HCoV-229E (like) viruses [Corman 2015]. Further evidence for this is the high diversity that HCoV-229E-like viruses detected in bats exhibit compared to those found in humans or camels, which suggests a long shared evolutionary history with bats.

The observed rather low host plasticity of HCoV-229E-like viruses among bats is noteworthy, since host plasticity has been proposed as a predictor for the likelihood of (successful) spillover into humans [Kreuder Johnson 2015]. However, in the absence of our knowledge of HCoV-229E-like viruses actually infecting camels and humans, one would potentially predict that these alpha CoVs might exhibit a lower risk for zoonotic spillover than more promiscuous bat CoVs such as Kenya bat coronavirus BtKY56 and Eidolon bat coronavirus/Kenya/KY24 [Kumakamba 2021]. In that respect there seem to be parallels between HCoV-229E and SARS-CoV-1 and SARS-CoV-2, which we believe to be primarily hosted by *Rhinolophus* bats [Li 2005; Lau 2005; Yip 2009; Yuan 2010; Hu 2017; Paraskevis 2020]. This counterintuitive observation might be the result of a sampling bias, but considering the comparably high number of sampled animals (2,581 bats in this study alone) it is one that can hardly be ignored. Surveillance and predictions of spillover risks particularly of bat CoVs will likely play an increasing role in the aftermath of the SARS-CoV-2 pandemic, but will face the challenge of limited data, despite the advances that have been made since the emergence of SARS-CoV-1.

While the zoonotic origin of globally circulating HCoV-229E is undisputed, the route by which it made its way from bats into humans is not clear. Some evidence, such as deletion patterns in the S gene and the open reading frames 4 and 8, suggests that camels may have served as an intermediate host. The hypothesis that an intermediate host was involved would be concurrent with what we know about the origins of other CoVs including as MERS-CoV (camel) or SARS-CoV-1 (civet) [Kan 2005; Corman 2014; Sabir 2016; Corman 2016; Forni 2017]. Phylogenetic evidence however, especially from the comparison of the more conserved genes, does not necessarily support that hypothesis. While the phylogenetic analyses of the most variable S- and the N-genes does place human and camel isolates into the same branch, this is not the case for the most conserved RdRp gene, and the E-, and M-genes (Figures 2-4, Supplement 4). This pattern would indicate that the transmission of HCoV-229E-like bat viruses into camels (which resulted in the isolates we know to date) may have been unrelated to the spillover that eventually led to the emergence of HCoV-229E. Such a multiple spillover scenario would not be without parallel, since CoVs closely related to SARS-CoV-2 were detected in pangolins, but overall clearly represent a different, earlier, and unrelated transmission [Zhang 2020; Lam 2020].

Determining the evolutionary history of HCoV-229E thus remains a challenge, not least due to limited and highly biased data available to date. Only 42 HCoV-229E isolates have been sequenced completely or near completely, with 28 derived from North America, 8 from Europe, and 7 from Southeast Asia - but not a single complete sequence from Africa. Similarly, almost all isolates from camels are derived from the Arabian peninsula and only one from Africa [Corman 2016; Li 2017].

### Infection rates are subject to seasonality

The detection of CoVs RNA in bats has been associated with seasonal differences in the past, and we found statistically significant evidence for such a correlation in our dataset as well [Anthony 2017; Montecino-Latorre 2020; Kumakamba 2021]. Much like in a recent study from the Democratic Republic of the Congo, we found that animals were more likely to be found shedding CoV RNA in the wet compared to the dry season, but that this is species dependent and may be true for some but be reversed for others (Table 3) [Kumakamba 2021]. Aside from this overall trend, the findings regarding seasonality from the two studies do not necessarily match up; however, this may be due to different bat species sampled on the one hand, and due to different ecological and climate conditions on the other hand. A potential reason for the seasonal differences may be related to the species’ birthing seasons, since it has been suggested that coronavirus transmission spikes in bat populations as juvenile bats become susceptible to infection once maternal antibody levels wane [Maganga 2020; Montecino-Latorre 2020]. The observation that young bats in our data set were more likely to be positive for CoV RNA than adults would support this hypothesis, though our finding was not statistically significant.

Interestingly, we found female bats to be significantly more likely to be CoV RNA positive than male bats, which is the opposite of what was found in DRC and what was suspected based on behavioral differences between males and females during breeding, birthing, and breastfeeding seasons (Kumakamba 2021; Fayenuwo 1974). However, while apparently a contradiction, this might be a reflection of differences in the species composition of the dataset, and indicate that caution should be used when making generalizations for members of the *Chiroptera* order, even if climate and habitat are similar.

### Closing remarks

Overall the results of this study on CoVs in African wildlife unveil or highlight several important aspects regarding the risks of future spillover and pandemics: 1) Our knowledge about the diversity of CoVs circulating in wildlife remains limited, as exemplified by the fact, that almost half of the CoV species we detected had not been described before. Despite having sampled 114 animal species, sample populations for most of them are too small to draw any conclusions about prevalence or risks. The role bats may play as a reservoir is certainly reinforced by the findings, but other species might simply be under sampled. The CoV RNA detection in a civet could be hinting towards a largely undetected CoV circulation in what could be an intermediate host for future spillovers. 2) The evidence for ecological factors such as seasonality driving transmission among bats is increasing. While the evidence remains circumstantial and the mechanisms elusive, it seems to become clear that further studies into this matter could enable smarter surveillance initiatives and mitigation measures such as limiting access to caves or discouraging hunting during periods of increased virus shedding. 3) The high prevalence and diversity of HCoV-229E-like viruses in African Hipposideridae bats reinforces the notion that HCoV-229E originated in Africa. However, while more and more data supports this origin hypothesis, the question whether or not camels acted as an intermediate host during the spillover into humans remains unclear. Regardless of the, or if there was an intermediate host, HCoV-229E highlights how important it is to not focus on Southeast Asia for CoV surveillance, but that CoV pandemics can start in many areas of the world.

## Supporting information

Legends for Supplements 3-5

## Acknowledgements

The authors would like to thank the Government of Cameroon for permission to conduct this study; the staff of Metabiota and CRESAR, who assisted in sample collection and testing; and any other involved members of the PREDICT-1 consortium (https://ohi.sf.ucdavis.edu/programs-projects/predict-project/authorship).

The study was undertaken as part of the global USAID-funded Emerging Pandemic Threats (EPT) PREDICT project, which focuses on enhancing the global capacity for the detection and discovery of potentially zoonotic viruses at the human-animal interface. It was made possible primarily by the generous support of the American people through the United States Agency for International Development (USAID) Emerging Pandemic Threats PREDICT program (cooperative agreement numbers GHN-A-OO-09-00010-00 and AID-OAA-A-14-00102). The contents are the responsibility of the authors and do not necessarily reflect the views of USAID or the United States Government. Some funding also came from the National Institutes of Health (NIH) Director’s Pioneer Award Program (grant number DP1-OD000370), the International Research Scientist Development Award from the NIH Fogarty International Center (K01 TW00003-1), Google.org, the Skoll foundation, the US Military HIV Research Program, and the Johns Hopkins Bloomberg School of Public Health, Center for a Livable Future.

## Statement of Data Availability

Additional data is available in the supplementary materials (Supplements 1-5). All viral sequences obtained are deposited in GenBank. Accession numbers for RdRp amplicons are listed in Table 2 and accession numbers for other partial genomic sequences of HCoV-229E-like viruses (MZ474745-MZ474805) are listed in Supplement 1.

