## Supplement 1 for "Wildlife in Cameroon harbor diverse coronaviruses including many isolates closely related to human coronavirus 229E"

**Supplement 1 Table: List of sequences obtained from CoVs related to HCoV-229E**

| Sample ID | RdRp | S | Aux | E | M | N | GenBank IDs |
| --- | --- | --- | --- | --- | --- | --- | --- |
| ECO06296 | part | part | - | - | part | - | KX284955; MZ474745; MZ474760 |
| CMAB71480 | part | part | full | full | full | part | MT081987; MZ474786 |
| CMAB71481 | part | part | - | - | part | part | MT082023; MZ474759; MZ474761; MZ474785 |
| CMAB71535 | part | part | full | full | part | - | MT221710; MZ474787 |
| CMAB71642 | part | part | full | full | full | part | MT081988; MZ474762; MZ474788 |
| CMAB71651 | part | part | full | full | full | part | MT081991; MZ474789 |
| CMAB71675 | part | part | full | full | full | part | MT082058; MZ474790 |
| CMAB71678 | part | part | - | - | part | - | MT082019; MZ474746; MZ474763 |
| CMAB71786 | part | part | - | - | - | - | MT082020; MZ474764 |
| CMAB71800 | part | part | - | - | part | - | MT082021; MZ474747; MZ474765 |
| CMAB71818 | part | part | full | full | full | part | MT081983; MZ474791 |
| CMAB72009 | part | part | full | full | full | part | MT081994; MZ474801 |
| CMAB72010 | part | part | full | full | part | - | MT082012; MZ474802 |
| CMAB72015 | part | part | full | full | part | part | MT081993; MZ474775; MZ474803 |
| CMAB72047 | part | part | - | - | - | - | MT081996; MZ474766 |
| CMAB72057 | part | part | - | - | part | - | MT082030; MZ474748; MZ474767 |
| CMAB72143 | part | part | full | full | part | part | MT082016; MZ474749; MZ474768; MZ474776; MZ474792 |
| CMAB72622 | part | part | - | - | part | - | MT063978; MT082053; MZ474750; MZ474769 |
| CMAB72642 | part | - | - | - | part | - | MT082018; MZ474751 |
| CMAB72694 | part | - | - | - | - | - | MT082026 |
| CMAB73354 | part | - | - | - | part | - | MT082057; MZ474752 |
| CMAB73355 | part | part | full | full | part | part | MT063996; MT082055; MZ474777; MZ474793 |
| CMAB73356 | part | - | - | - | - | part | MT082031; MZ474778 |
| CMAB73357 | part | part | part | part | full | part | MT063997; MT082056; MZ474794; MZ474795 |
| CMAB73364 | part | part | full | full | part | part | MT063985; MT082006; MZ474779; MZ474796 |
| CMAB73367 | part | - | - | - | - | - | MT082029 |
| CMAB73368O | part | - | - | - | - | - | MT064018 |
| CMAB73368R | part | part | full | full | part | part | MT063980; MT082008; MZ474753; MZ474780; MZ474797 |
| CMAB73372 | part | part | - | - | part | part | MT063984; MT082010; MZ474754; MZ474770; MZ474781 |
| CMAB73376 | part | part | full | full | full | part | MT064047; MT082009; MZ474798 |
| CMAB73401 | part | - | - | - | part | part | MT063999; MZ474755; MZ474782 |
| CMAB74746 | part | - | - | - | - | - | MT064013; MT082084 |
| CMAB74747 | part | - | - | - | - | - | MT064014 |
| CMAB74760 | part | - | - | - | - | - | MT082078 |
| CMAB75005 | part | part | - | - | - | - | MT064022; MT082090; MZ474771 |
| CMAB75013O | part | part | part | part | part | - | MT064024; MT082091; MZ474799; MZ474804 |
| CMAB75013R | part | part | part | part | part | - | MT064025; MT082092; MZ474783; MZ474800; MZ474805 |
| CMAB75014 | part | part | - | - | part | part | MT064026; MT082093; MZ474756; MZ474772; MZ474784 |
| CMAB75019 | part | part | - | - | part | - | MT064028; MT082081; MZ474757; MZ474773 |
| CMAB75026 | part | part | part | part | part | - | MT064029; MZ474758; MZ474774 |
| CMAB75030 | part | - | - | - | - | - | MT064031 |
| CMAB75051 | part | - | - | - | - | - | MT064035 |
