## Supplement 2 for "Wildlife in Cameroon harbor diverse coronaviruses including many isolates closely related to human coronavirus 229E"

**Supplement 2 Table: List of animal species sampled**

| ***Bats (n=2581)*** |  |
| --- | --- |
| *Casinycteris argynnis* | *2* |
| *Chaerephon major* | *1* |
| *Chaerephon pumilus* | *31* |
| *Chiroptera* (unspecified species) | *1* |
| *Coleura afra* | *2* |
| *Eidolon helvum* | *267* |
| *Epomophorus gambianus* | *32* |
| *Epomops franqueti* | *206* |
| *Glauconycteris poensis* | *2* |
| *Hipposideros beatus* | *2* |
| *Hipposideros caffer* | *165* |
| *Hipposideros curtus* | *7* |
| *Doryrhina cyclops* | *37* |
| *Hipposideros fuliginosus* | *6* |
| *Macronycteris gigas* | *163* |
| *Hipposideros ruber* | *653* |
| *Hipposideros sp.* | *2* |
| *Hypsignathus monstrosus* | *7* |
| *Hypsugo musciculus* | *6* |
| *Kerivoula cuprosa* | *5* |
| *Kerivoula sp.* | *2* |
| *Lavia frons* | *3* |
| *Lissonycteris angolensis* | *62* |
| *Megaloglossus woermanni* | *184* |
| *Micropteropus pusillus* | *137* |
| *Miniopterus inflatus* | *2* |
| *Miniopterus schreibersii* | *1* |
| *Molossidae* | *1* |
| *Mops condylurus* | *175* |
| *Mops demonstrator* | *7* |
| *Myonycteris torquata* | *37* |
| *Nanonycteris veldkampii* | *3* |
| *Neoromicia capensis* | *3* |
| *Neoromicia nana* | *2* |
| *Neoromicia tenuipinnis* | *13* |
| *Nycteris grandis* | *23* |
| *Nycteris hispida* | *23* |
| *Nycteris major* | *1* |
| *Nycteris thebaica* | *1* |
| *Nycticeinops schlieffenii* | *3* |
| *Pipistrellus inexspectatus* | *4* |
| *Pipistrellus nanulus* | *8* |
| *Pipistrellus rusticus* | *2* |
| *Rhinolophus alcyone* | *4* |
| *Rhinolophus cf. alcyone* | *1* |
| *Rhinolophus fumigatus* | *1* |
| *Rhinolophus landeri* | *19* |
| *Rhinopoma microphyllum* | *1* |
| *Rousettus aegyptiacus* | *200* |
| *Scotoecus hirundo* | *1* |
| *Scotonycteris zenkeri* | *6* |
| *Scotophilus dinganii* | *12* |
| *Scotophilus leucogaster* | *33* |
| *Scotophilus nux* | *3* |
| *Taphozous mauritianus* | *4* |
| *Vespertilionidae* | *2* |
| ***Carnivores (n=37)*** |  |
| *Civettictis civetta* | *8* |
| *Herpestes naso* | *4* |
| *Nandinia binotata* | *25* |
| ***Eulipotyphla (n=159)*** |  |
| *Crocidura goliath* | *2* |
| *Crocidura poensis* | *36* |
| *Crocidura sp.* | *4* |
| *Soricidae (unidentified genus)* | *64* |
| *Sylvisorex sp.* | *53* |
| ***Primates (n=1006)*** |  |
| *Cercocebus agilis* | *3* |
| *Cercocebus torquatus* | *5* |
| *Cercopithecus cephus* | *531* |
| *Cercopithecus erythrotis* | *2* |
| *Cercopithecus mona* | *5* |
| *Cercopithecus neglectus* | *26* |
| *Cercopithecus nictitans* | *228* |
| *Cercopithecus pogonias* | *5* |
| *Cercopithecus preussi* | *2* |
| *Cercopithecus wolfi* | *57* |
| *Chlorocebus aethiops* | *9* |
| *Chlorocebus tantalus* | *2* |
| *Colobus guereza* | *22* |
| *Colobus satanas* | *3* |
| *Erythrocebus patas* | *1* |
| *Gorilla gorilla* | *5* |
| *Mandrillus leucophaeus* | *8* |
| *Mandrillus sphinx* | *13* |
| *Miopithecus talapoin* | *7* |
| *Pan troglodytes* | *40* |
| *Papio anubis* | *9* |
| *Papio hamadryas* | *22* |
| *Perodicticus potto* | *1* |
| ***Hyraxes (n=2)*** |  |
| *Dendrohyrax dorsalis* | *2* |
| ***Pangolins (n=38)*** |  |
| *Phataginus tricuspis* | *38* |
| ***Rodents (n=2740)*** |  |
| *Acomys johannis* | *3* |
| *Arvicanthis sp.* | *1* |
| *Atherurus africanus* | *1048* |
| *Cricetomys emini* | *64* |
| *Dasymys rufulus* | *12* |
| *Deomys ferrugineus* | *6* |
| *Dephomys sp.* | *1* |
| *Funisciurus isabella* | *3* |
| *Gerbilliscus kempi* | *10* |
| *Heimyscus fumosus* | *2* |
| *Heliosciurus rufobrachium* | *1* |
| *Hybomys univittatus* | *3* |
| *Hylomyscus sp.* | *7* |
| *Lemniscomys striatus* | *20* |
| *Lemniscomys sp.* | *4* |
| *Lophuromys nudicaudus* | *115* |
| *Lophuromys sikapusi* | *6* |
| *Malacomys longipes* | *7* |
| *Malacomys sp.* | *6* |
| *Mastomys sp.* | *19* |
| *Muridae (unidentified genus)* | *10* |
| *Mus musculus* | *180* |
| *Mus setulosus* | *605* |
| *Mus sp.* | *96* |
| *Oenomys hypoxanthus* | *17* |
| *Praomys jacksoni* | *71* |
| *Praomys misonnei* | *34* |
| *Praomys sp.* | *131* |
| *Protoxerus stangeri* | *1* |
| *Rattus rattus* | *101* |
| *Rodentia (unidentified family)* | *7* |
| *Stochomys longicaudatus* | *7* |
| *Thryonomys swinderianus* | *140* |
| *Uranomys ruddi* | *1* |
| *Xerus erythropus* | *1* |
| ***Even-toed ungulates (n=17)*** |  |
| *Cephalophus callipygus* | *3* |
| *Cephalophus dorsalis* | *6* |
| *Philantomba monticola* | *7* |
| *Potamochoerus porcus* | *1* |
