## Supplementary figures and images for "Wildlife in Cameroon harbor diverse coronaviruses including many isolates closely related to human coronavirus 229E"

### Supplement 3

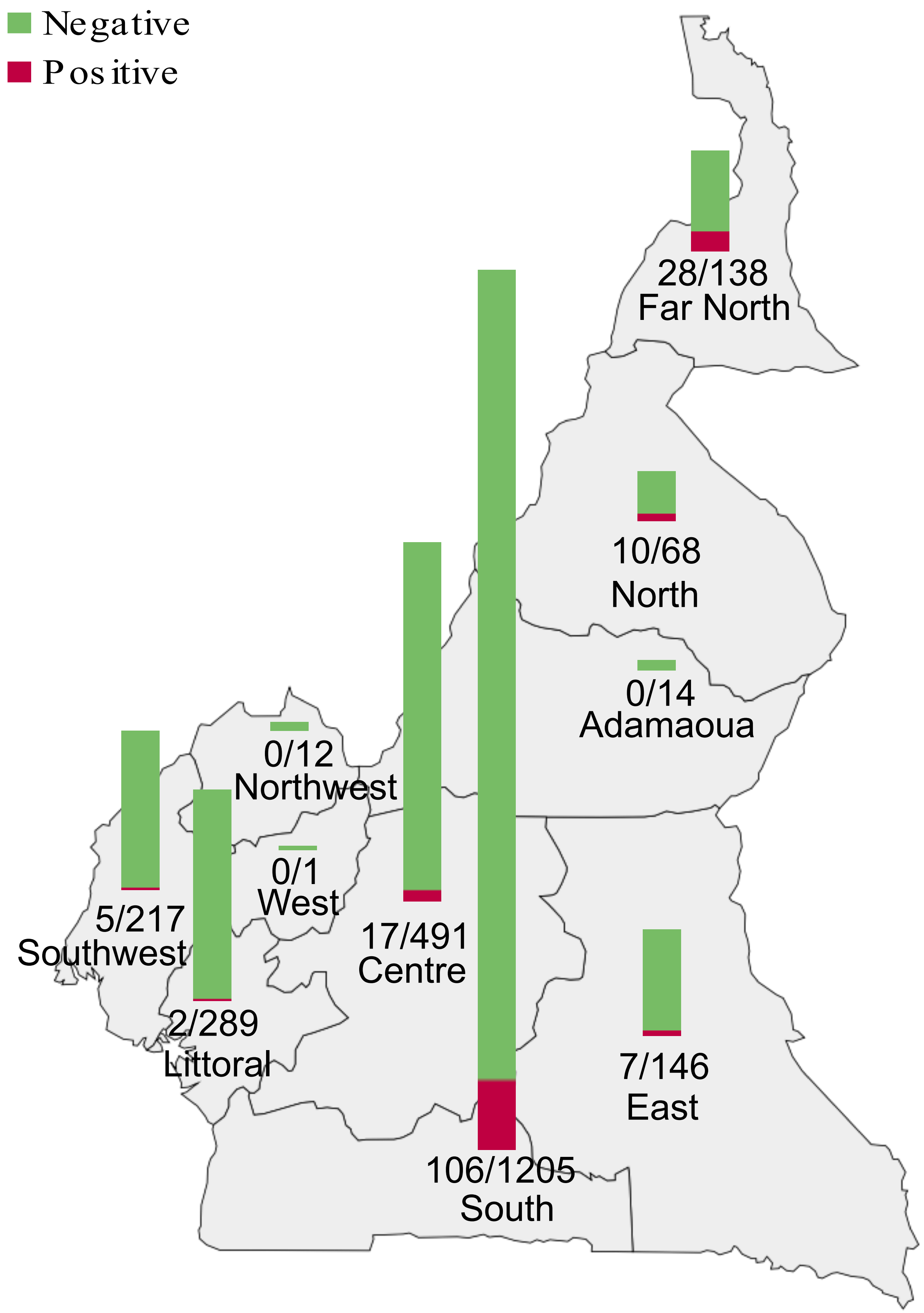

### Supplement 4

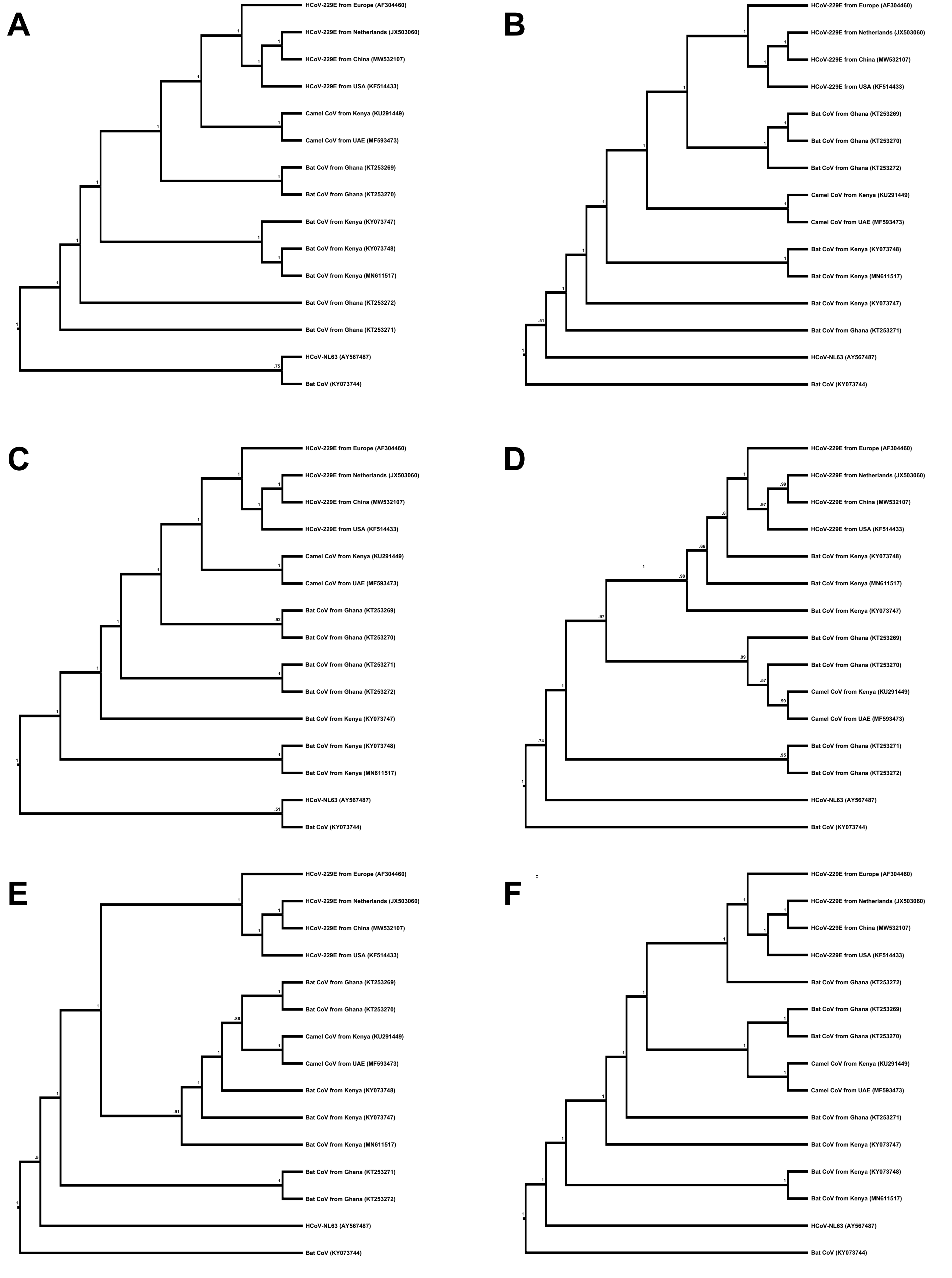

### Supplement 5

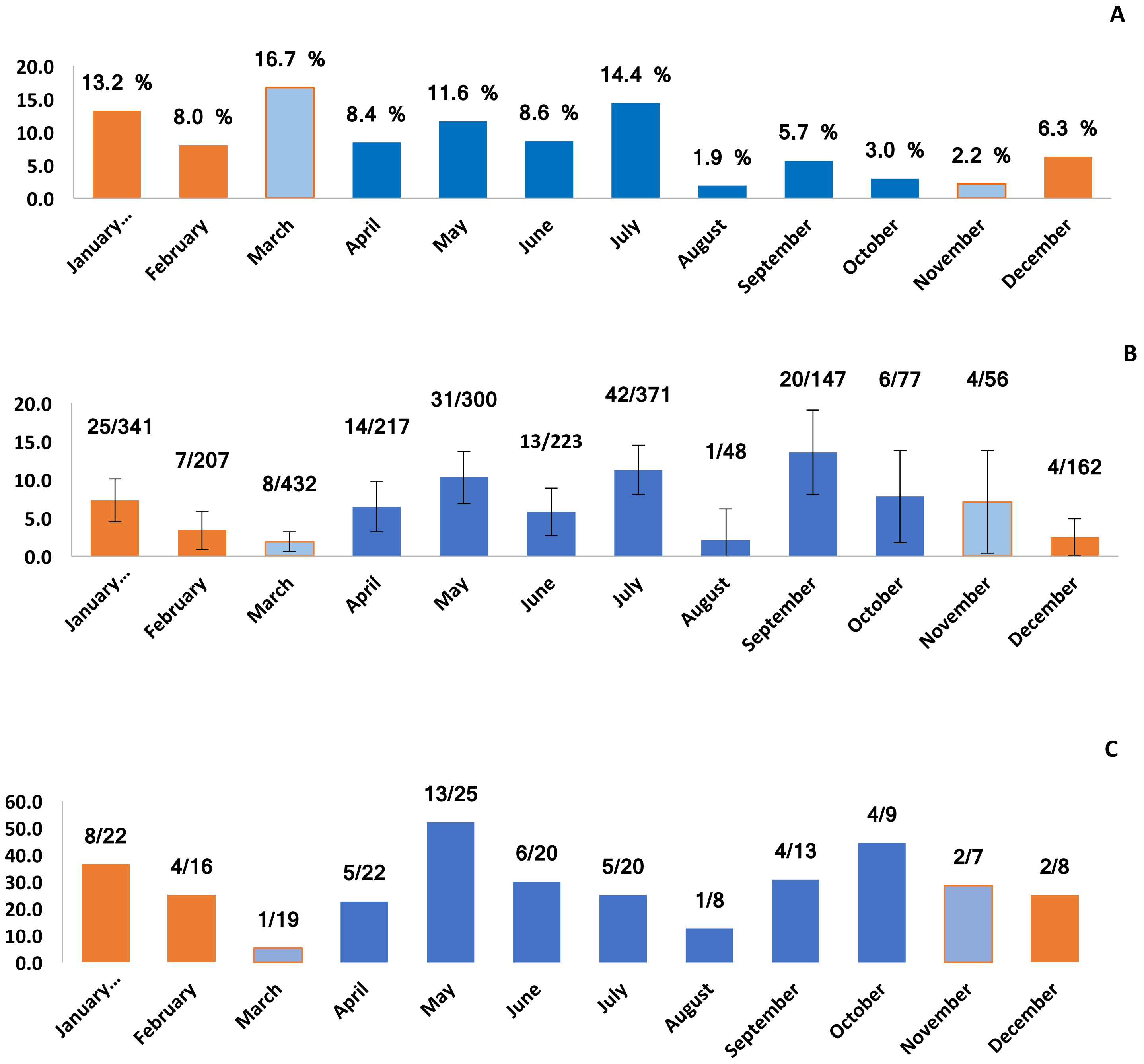
