## Supplementary material for "Wildlife in Cameroon harbor diverse coronaviruses including many isolates closely related to human coronavirus 229E": Legends for Supplements 3-5

**Figure legends, Supplementary Figures:**

**Supplement 3 Figure: Sampling effort by region (bats only)**

**Supplement 4 Figure, Phylogenetic trees of HCoC-229E:** Maximum likelihood phylogenetic trees of coronavirus sequences related to HCoC-229E, based on the Full genome (A), RdRp (B), Spike (C), Envelope (D), Membrane (E) and Nucleoprotein (F). Numbers at nodes indicate bootstrap support.

**Supplement 5 Figure, Bat sampling effort:** Bat sampling effort by month indicating proportion of total samples (A), proportion of individual animals positive for coronavirus RNA (B) and proportion of sampling events with at least one bat positive for coronavirus RNA.
